# Dietary- and host-derived metabolites are used by diverse gut bacteria for anaerobic respiration

**DOI:** 10.1101/2022.12.26.521950

**Authors:** Alexander S. Little, Isaac T. Younker, Matthew S. Schechter, Paola Nol Bernardino, Raphaël Méheust, Joshua Stemczynski, Kaylie Scorza, Michael W. Mullowney, Deepti Sharan, Emily Waligurski, Rita Smith, Ramanujam Ramanswamy, William Leiter, David Moran, Mary McMillin, Matthew A. Odenwald, Anthony T. Iavarone, Ashley M. Sidebottom, Anitha Sundararajan, Eric G. Pamer, Murat A. Eren, Samuel H. Light

**Author notes:** Address correspondence to Samuel H. Light.

## Abstract

Respiratory reductases enable microbes to utilize molecules present in anaerobic ecosystems as energy-generating respiratory electron acceptors. Here we identify three taxonomically distinct families of human gut bacteria (Burkholderiaceae, Eggerthellaceae, Erysipelotrichaceae) that encode large arsenals of tens-to-hundreds of respiratory-like reductases per genome. Screening species from each family (*Sutterella wadsworthensis*, *Eggerthella lenta*, and *Holdemania filiformis*), we discover 22 metabolites used as respiratory electron acceptors in a species-specific manner. Identified reactions transform multiple classes of dietary- and host-derived metabolites, including bioactive molecules resveratrol and itaconate. Products of identified respiratory metabolisms highlight poorly characterized compounds, such as the itaconate-derived 2-methylsuccinate. Reductase substrate-profiling defines enzyme-substrate pairs and reveals a complex picture of reductase evolution, providing evidence that reductases with specificities for related cinnamate substrates independently emerged at least four times. These studies thus establish an exceptionally versatile form of anaerobic respiration that directly links microbial energy metabolism to the gut metabolome.

## Introduction

Heterotrophic cellular respiration is defined by the oxidation of an organic electron donor and the passage of resulting electrons through an electron transport chain to a terminal electron acceptor.^1^ Electron transfer is coupled to the creation of an ion gradient which, in turn, powers oxidative adenosine triphosphate (ATP) synthesis through ATP synthase.^1^ While molecular oxygen is a hallmark respiratory electron acceptor, microbes residing in oxygen-poor environments possess respiratory metabolisms that utilize alternative electron acceptors.^1^

Fermentative metabolisms dominate the anaerobic gut microbial ecosystem, but several classical respiratory metabolisms have been identified. Diverse sulfate-reducing bacteria use different sulfate respiratory electron acceptors.^2^ Clostridial acetogens and Archaeal methanogens respire carbon dioxide.^3^ Bacteroidales convert fermentative substrates to the electron acceptor fumarate, which is reduced by the enzyme fumarate reductase.^4^ Various other gut bacteria convert exogenous 4-carbon dicarboxylates, including the amino acid aspartate, to fumarate, which is similarly respired.^5^

In the inflamed gut, electron acceptors generated by immune cells (hydrogen peroxide, tetrathionate, and nitrate) fuel respiratory growth of pathogenic and commensal Enterobacteriaceae.^6–8^

While most identified respiratory gut metabolisms use inorganic electron acceptors, two observations suggest that other types of respiratory electron acceptors may be underappreciated. First, a recent study found that *Eggerthella lenta* used the neurotransmitter dopamine as a respiratory electron acceptor and identified several related catechols that were similarly reductively dexhydroxylated.^9^ Second, a bioinformatic survey found that multiple Eggerthellaceae family bacteria encode many enzymes homologous to the respiratory dimethylsulfoxide (DMSO) reductase and noted that they might enable use of different electron acceptors.^10^ These observations underscore the possibility that gut microbes possess the capacity to utilize a large and chemically diverse collection of respiratory electron acceptors.

Here we employ a genome mining-based approach to interrogate respiratory electron acceptor usage in the human gut microbiome. We find that three taxonomically distinct families of gut microbes encode exceptionally large numbers of respiratory-like reductases. We show that representative species from each clade exhibit respiratory activities and identify diverse small molecule electron acceptors used in a strain-dependent manner. We additionally pair specific reductases with their substrates to reveal a convoluted relationship between reductase sequence and substrate specificity. By probing reductases with distinct active site architectures, we show that parallel evolutionary trajectories generated reductases that achieve similar activities through distinct mechanisms. These studies establish a distinctive mode of bacterial respiration, defined by the versatile use of organic electron acceptors, and contextualize the role of energy metabolism in shaping the gut metabolome.

## Results

### Distantly related bacteria encode large reductase arsenals

Previous studies have identified a number of microbial respiratory reductases that enable use of distinct electron acceptors. Many of these reductases are evolutionarily related and placed within one of two enzyme superfamilies by hidden Markov models (HMM). Members of the “molybdopterin” superfamily (HMM PF00384) – exemplified by dimethylsulfoxide (DMSO) reductase – contain a catalytic domain that binds the redox-active cofactor molybdopterin and tend to act on inorganic respiratory electron acceptors (**Fig. 1A, Supplementary Table 1**).^9,11–22^ Members of the “flavin” superfamily (HMM PF00890) – exemplified by fumarate and urocanate reductases – contain a catalytic domain that binds the redox-active cofactor flavin adenine dinucleotide and tend to act on organic respiratory electron acceptors (**Fig. 1B, Supplementary Table 1**).^23–28^ Structural and biochemical studies of enzymes within the molybdopterin and flavin superfamilies suggest each contains a broadly conserved electron transfer mechanism but active site distinctions that confer different substrate specificities and activities.^29,30^

**Figure 1.**
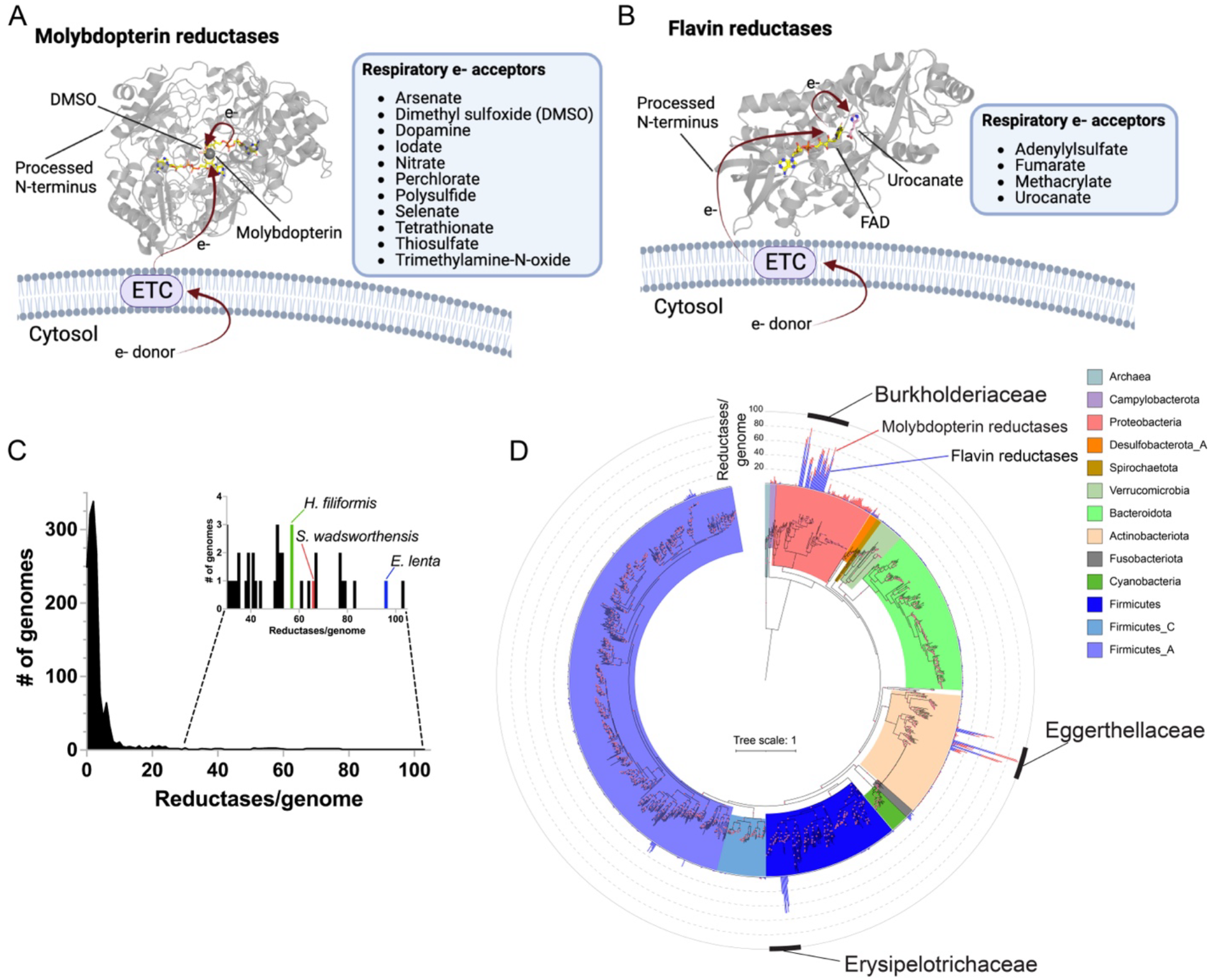
Respiratory reductase orthologs are highly overrepresented in three distinct lineages of gut bacteria. (A) General mechanism and electron acceptors used by different previously characterized respiratory molybdopterin reductases (Pfam PF00384). Arrows highlight electron (e^-^) transfer path from the electron transport chain (ETC) to dimethylsulfoxide (DMSO) in the DMSO reductase crystal structure (PDB code 4DMR). Electron acceptors and general electron transfer mechanism used by previously characterized respiratory flavin reductases (Pfam PF00890). Arrows highlight electron transfer to substrate in the *S. oneidensis* urocanate reductase co-complex crystal structure (PDB code 6T87). (C) Distribution of the number of flavin/molybdopterin reductases in 1533 representative human gut bacteria genomes and metagenome-assembled genomes. (D) Phylogenetic reconstruction of the evolutionary history of genomes analyzed in (C). The maximum likelihood tree was constructed based on a concatenated alignment of 14 ribosomal proteins under an LG + I + G4 model of evolution (2092 amino acid sites). The number of flavin (blue) and molybdopterin (red) reductases with a computationally predicted signal peptide in each genome are graphed on the outer ring of the tree.

Most characterized respiratory reductases are soluble proteins that receive electrons from the electron transport chain in the cytosolic membrane. While respiratory reductases can be associated with either the cytosolic or extracytosolic side of the cytosolic membrane, we observed most characterized flavin and molybdopterin respiratory reductases contain an N-terminal signal peptide characteristic of extracytosolic localization (**Supplementary Table 1**). By contrast, most molybdopterin and flavin superfamily enzymes that possess an obvious non-respiratory reductase functionality are cytosolically localized and lack an N-terminal signal peptide (**Supplementary Table 1**).

Based on the role of established flavin and molybdopterin reductases in respiratory electron acceptor usage, we reasoned that a survey of genes predicted to encode secreted flavin or molybdopterin superfamily members could facilitate the identification of respiratory gut microbes. We analyzed 1533 genomes and metagenome-assembled genomes from a recently complied collection of diverse human gut prokaryotes and observed a striking distribution in the number of reductases per genome.^31^ While most genomes encode less than 5 reductases, we identified a small subset of ‘high reductase’ genomes that encode >30 (and as many as 103) reductases (**Fig. 1C, Supplementary Table 2**). Consistent with identified reductases having an extracellular localization and respiratory functionality, most contained a predicted N-terminal signal peptide (**Supplementary Table 2**).

Further taxonomic analyses revealed ‘high reductase’ genomes form three distinct clades that span multiple genera in the: (1) Actinobacteria family Eggerthellaceae, (2) Firmicutes family Erysipelotrichaceae, and (3) Proteobacteria family Burkholderiaceae (**Fig. 1D**). Notably, species from each clade – including *E. lenta* from Eggerthellaceae, *Sutterella wadsworthensis* from Burkholderiaceae, and *Holdemania filiformis* from Erysipelotrichaceae – are prevalent, low-to-moderate abundance members of the human gut microbiome.^32^ Expanding the analyses to include bacteria found outside the human gut revealed that high reductase clades include >200 reductases/genome bacteria and exhibit complex reductase gain/loss patterns indicative of a convoluted evolutionary history (**Extended Data Fig. 1, Supplementary Table 3, Supplementary Discussion**). These analyses thus define three taxonomically distinct groups of gut bacteria that encode large numbers of respiratory-like reductases.

### Reductase-encoding bacteria exhibit respiratory growth

To test whether gut microbes with a high number of reductases per genome possessed respiratory capabilities, we selected *E. lenta*, *S. wadsworthensis*, and *H. filiformis* strains for experimental characterization. We identified a flavin reductase with >50% sequence identity to a previously characterized respiratory urocanate reductase (UrdA) encoded within each strain’s genome, suggesting that the small molecule urocanate provided a suitable candidate for preliminary investigations of respiratory growth.^24,25^

As use of a respiratory electron acceptor is conditional upon oxidation of an electron donor, we screened common respiratory electron donors for the ability to induce urocanate-dependent growth enhancement. We observed a synergistic formate/urocanate-dependent growth enhancement of *E. lenta* and *S. wadsworthensis*, establishing formate as a viable electron donor for these strains (**Fig. 2A**). *H. filiformis* also exhibited enhanced growth in the presence of urocanate, but requirements for a rich growth medium hindered identification of a respiratory electron donor (**Fig. 2A**). Consistent with the observed phenotypes reflecting respiratory activity, growth enhancement of each strain tracked with urocanate reduction to imidazole propionate and depended upon the electron-accepting properties of urocanate (i.e., the reduced reaction product did not enhance growth) (**Fig. 2A-2C**). Further supporting the role of respiration in these growth phenotypes, we found that urocanate stimulated ATP synthesis of all three strains (**Fig. 2D**). These results thus demonstrate that *E. lenta*, *S. wadsworthensis*, and *H. filiformis* possess respiratory metabolic capabilities and similarly utilize urocanate as a respiratory electron acceptor.

**Figure 2.**
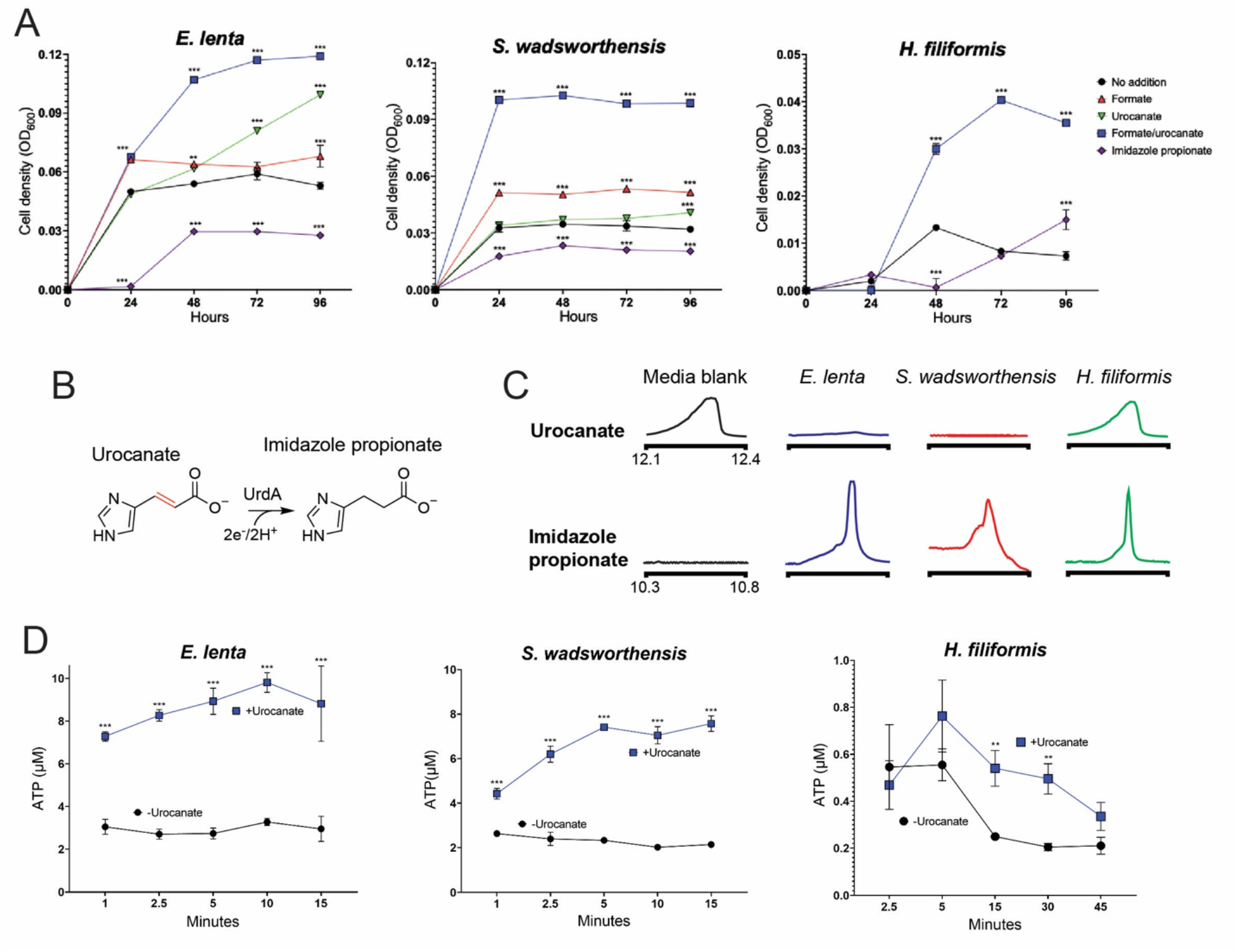
‘High reductase’ gut bacteria exhibit respiratory growth properties. (A) Growth of *E. lenta* DSM2243, *S. wadsworthensis* DFI.4.78, and *H. filiformis* DSM12042 strains in the presence of electron donor (formate), electron acceptor (urocanate), and product of urocanate reduction (imidazole propionate). (B) Reaction catalyzed by urocanate reductase UrdA. (C) GC-MS extracted ion chromatograms in formate/urocanate-supplemented culture supernatants following cultivation of the indicated strains. (D) ATP produced by cells suspended in formate-supplemented buffer. For (A) data are mean ±SD (n = 3 independent biological replicates), for (D) data are mean ±SD (n = 3 technical replicates). *p < 0.05, **p < 0.01, *** p < 0.001. Two-way ANOVA, multiple test vs media alone.

### Reductase-encoding bacteria use diverse electron acceptors

Having established the respiratory capabilities of our strains, we hypothesized that their large number of reductases reflected an extreme versatility in respiratory electron acceptor utilization capabilities. Since most reductases encoded by these strains were distantly related to functionally characterized enzymes and thus lacked obvious substrates, we assembled a panel of metabolites that possessed electron-accepting properties and could be present in the gastrointestinal tract.

By screening *E. lenta*, *S. wadsworthensis* and *H. filiformis* stains, we identified nineteen compounds that supported growth of at least one species (**Fig. 3A, Extended Data Figs. 2-4**). For most identified compounds we confirmed that: (1) their depletion coincided with accumulation of a reduced product, (2) this reduced product did not impact growth, and (3) presence of an electron donor was required for growth enhancement (**Extended Data Figs. 2-4**). Through complementary enzymatic activity assays, we identified three additional molecules (resveratrol, catechin, and epicatechin) that were used as electron acceptors but, due to poor solubility or other factors, were not associated with enhanced growth under the tested conditions **(Fig. 3B & 3C, Extended Data Fig. 5**). Further corroborating the respiratory role of observed activities, we confirmed the representatives from each class of identified electron acceptor promoted ATP production (**Extended Data Fig. 6**).

**Figure 3.**
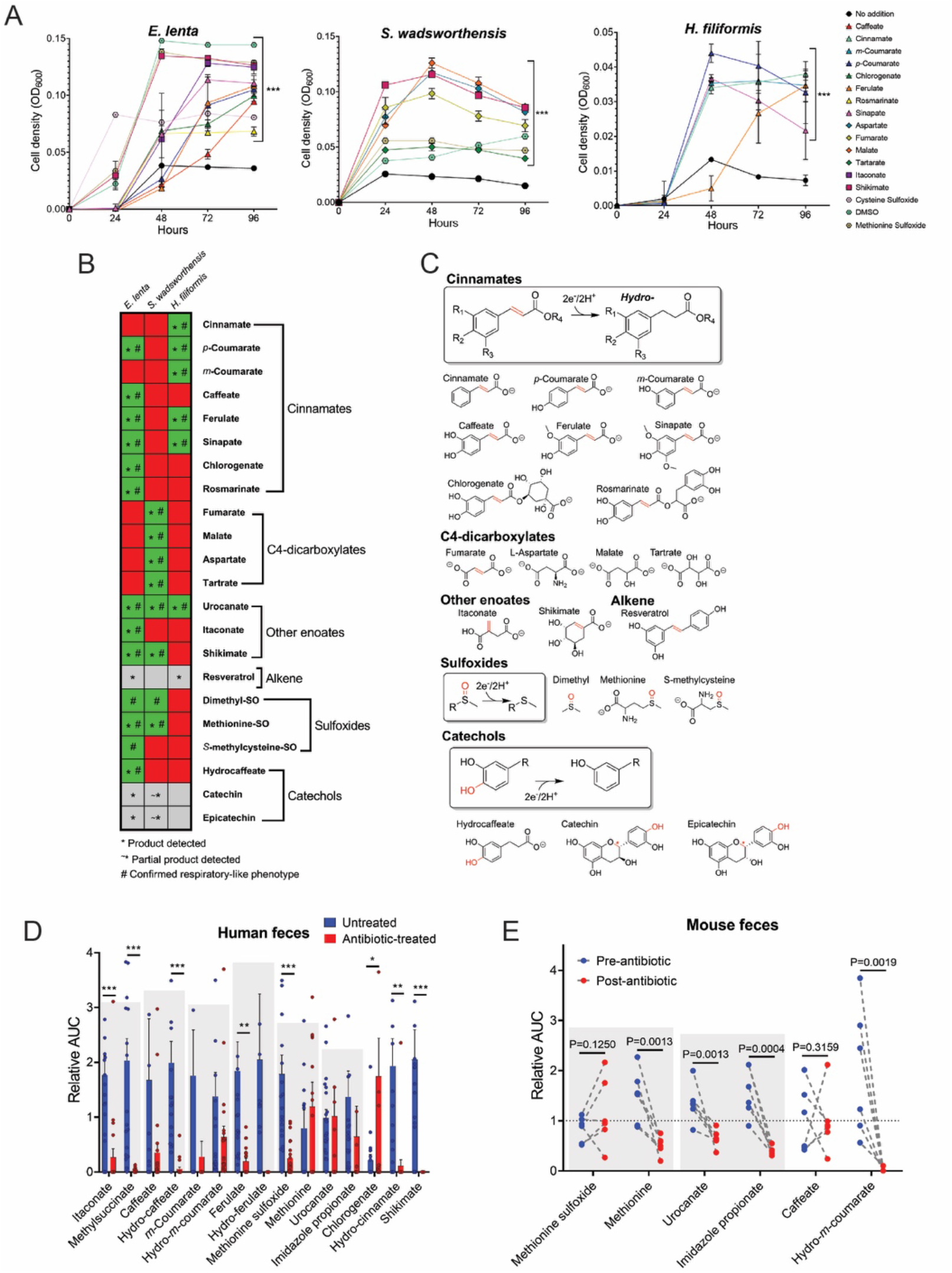
‘High reductase’ gut bacteria utilize diverse respiratory electron acceptors. (A) *E. lenta* DSM2243, *S. wadsworthensis* DFI.4.78, and *H. filiformis* DFI.9.20 growth in media supplemented with formate and identified growth-stimulating small molecules. (B) Summary of electron acceptor usage findings. Green indicates growth enhancement, red indicates no enhancement, and grey denotes limited solubility. The symbol * indicates the reduced product was detected after growth, and the symbol # indicates a confirmed respiratory phenotype. (C) Summary of identified reductase activities. The electron-accepting bond is highlighted in red. The red symbol * for both catechin and epicatechin denotes the site of benzylic C-O cleavage. (D) Identified metabolites in fecal samples collected from non-antibiotic versus antibiotic treated subjects as measured by LC-MS. Area under the curve (AUC) values are presented relative to each metabolite’s normalized average (normalized to 1) within the dataset as measured by LC-MS. See **Supplementary Table 4** for original data. (E) Identified metabolites in mouse feces pre- and post-antibiotic treatment. Area under the curve (AUC) values are presented relative to each metabolite’s normalized average (normalized to 1) within the dataset. Gray boxes highlight reductase substrate and product pairs. See **Supplementary Table 5** for original data. For (A) data are mean ±SD (n = 3 independent biological replicates) with two-way ANOVA, multiple test vs media alone. For (D) data are mean ±SEM (n = 21 antibiotic treated, 20 untreated) with multiple unpaired t tests. For (E) n = 6 pre- and 6-post antibiotic treated, with multiple unpaired t tests. *p < 0.05, **p < 0.01, *** p < 0.001.

Identified electron acceptors included multiple cinnamates, flavonoids, 4-carbon-dicarboxylates, sulfoxides, with electron-accepting groups including carbon-oxygen bonds, hydroxyls, sulfoxides, and alkenes (**Fig. 3A-3C**). Different substrate classes exhibited species-specific patterns of utilization (further explored in **Supplementary Discussion**). In addition to these broader substrate classes, we found that (1) *E. lenta* and *S. wadsworthensis* reduced an enoate group in shikimate, an intermediate in the biosynthesis of aromatic compounds (**Fig. 3A-C, Extended Data Figs. 2 & 3**), (2) *E. lenta* and *H. filiformis* reduced an alkene group in resveratrol, a plant phenylpropanoid present in different foods/beverages that has many reported effects on mammalian physiology (**Extended Data Fig. 5**) and (3) *E. lenta* reduced an alkene group in itaconate, a mammalian immune signaling molecule (**Fig. 3C, Extended Data Fig. 2**).^33–35^ Collectively, these results define a complex species-dependent pattern of electron acceptor usage.

### Reductase substrates and products are present in feces

To clarify the relevance of identified electron acceptors within the gut microbiome, we measured fecal metabolites from a population of 19 non-antibiotic-treated healthy donors and 22 hospitalized antibiotic-treated patients with severely diminished microbiome complexity (**Extended Data Fig. 8A**). As the multiple differences that distinguish these healthy and antibiotic-treated groups precludes the definitive attribution of metabolite differences to the microbiome, we additionally sought a more controlled experimental system. We thus measured electron acceptors and their reduced products in mouse fecal samples before and after treatment with a cocktail of broad-spectrum antibiotics that severely disrupted the gut microbiome (**Extended Data Fig. 8B**).

Distinct compounds were detected in the mice and human samples but, in both populations, levels of reductase substrates and products trended lower in antibiotic-treated groups (**Fig. 3D & 3E, Supplementary Tables 4 & 5**). This pattern was evident for multiple cinnamates, which can be produced from bacterial breakdown of larger dietary polyphenols, urocanate and shikimate, which are intermediates in bacterial metabolic pathways and, perhaps most strikingly, for the immunometabolite itaconate (**Fig. 3D, Supplementary Table 4**). Itaconate and its reduced product 2-methylsuccinate were detected in virtually every healthy human fecal sample but were below the limit of detection for most samples from the antibiotic-treated group (**Fig. 3D, Supplementary Table 4**). These studies establish that the identified respiratory electron acceptors are present within the mammalian gastrointestinal tract and responsive to the gut microbiome.

### Respiratory electron acceptors selectively induce reductases

An examination of the genomic context of *E. lenta*, *S. wadsworthensis*, and *H. filiformis* flavin and molybdopterin reductase genes revealed that they frequently colocalized with putative transcriptional regulators, though notable species-specific differences in the type of regulators were evident. *S. wadsworthensis* and *H. filiformis* reductase genes often colocalized with histidine kinase two-component systems or typical helix-turn-helix cytosolic transcriptional regulators, whereas 64 *E. lenta* reductase genes directly neighbor a recently described family of transmembrane transcriptional regulators (**Fig. 4A & 4B, Supplementary Table 6**).^36^ As bacterial transcriptional regulators often establish autoregulatory circuits that concordantly regulate neighboring genes on the genome, we reasoned it might be possible to leverage transcriptional responses to respiratory electron acceptors to gain insight into reductase substrate specificity.

**Figure 4.**
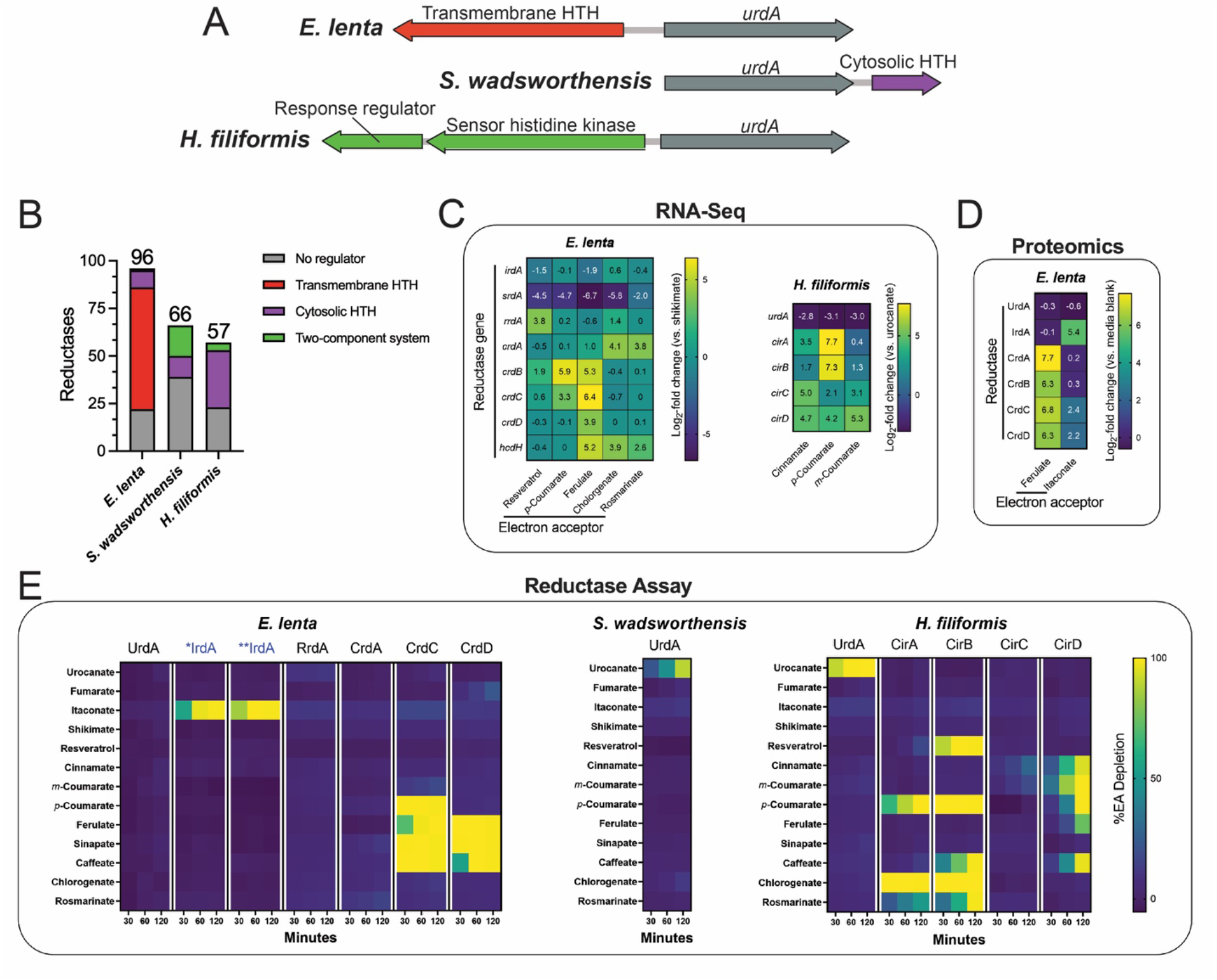
Flavin reductases are induced by their electron acceptors and exhibit relatively narrow substrate specificities. (A) Genomic context of *urdA* urocanate reductases in *E. lenta* DSM2243, *S. wadsworthensis* DFI.4.78, and *H. filiformis* DFI.9.20 genomes. Genes encoding predicted transcriptional regulators are noted. (B) Genomic context of flavin and molybdopterin reductases with respect to adjacent regulatory elements. Either no clear element present (grey), two-component system (green), cytosolic regulator (purple), or transmembrane regulator (red). Cytosolic and transmembrane refers to the cellular localization of the putative signal-receiving domain in predicted transcriptional regulators that contain a DNA-binding helix-turn-helix (HTH) domain. The number of flavin and molybdopterin reductases genes that directly neighbor a predicted transcriptional regulator in *E. lenta* DSM2243, *S. wadsworthensis* DFI.4.78, and *H. filiformis* DFI.9.20 genomes. RNA-Seq results from *E. lenta* DSM2243 and *H. filiformis* DFI.9.20 cells cultivated in media supplemented with indicated electron acceptors. (D) Proteomics results from *E. lenta* DSM2243 cells cultivated in media supplemented with indicated electron acceptors. (E) Activity of recombinant reductases in the presence of indicated electron acceptors. For (E), representative data are shown.

After confirming the predicted *E. lenta* urocanate reductase was one of the most strongly induced genes in the presence of urocanate, we performed RNA-seq on *E. lenta* and *H. filiformis* in nine conditions containing a respiratory electron acceptor used by an unknown reductase (**Fig. 4C, Supplementary Tables 7 & 8**). We further performed proteomic analyses of *E. lenta* grown in the presence of ferulate and itaconate to identify reductase induction at the protein level (**Fig. 4D, Supplementary Table 9**). From these experiments, we identified at least one flavin reductase as among the most strongly induced genes in the presence of each electron acceptor. To facilitate downstream analyses, we assigned reductase gene names based on the condition they were induced (*crdA* for the *E. lenta* cinnamate-induced reductase A, *cirB* for the *H. filiformis* cinnamate-induced reductase B, etc.) (**Supplementary Table 10**).

We next compared reductase expression across conditions and observed a complex pattern of regulation. While itaconate, shikimate, and resveratrol each specifically induced a single flavin reductase, distinct cinnamates differentially induced multiple flavin reductases (**Fig. 4C & 4D**). For both *E. lenta* and *H. filiformis,* structurally related cinnamates resulted in elevated but variable expression of four distinct flavin reductases (**Fig. 4C**). Furthermore, in *E. lenta*, ferulate and chlorogenate, but not *p*-coumarate, induced the previously characterized hydrocaffeate dehydroxylase *hcdh* (**Fig. 4C**).^9^ These findings thus reveal specific but sometimes complex regulatory responses of reductases to related respiratory electron acceptors.

### Reductase induction predicts substrate specificity

To assess whether gene expression patterns could predict enzyme substrate specificity, we recombinantly expressed thirteen flavin reductases identified in our transcriptomics/proteomics studies (**Extended Data Fig. 9**). Due to poor yields of the *E. lenta* itaconate-induced reductase, IrdA, we additionally expressed two close IrdA orthologs from *Adlercreutzia muris* and *Berryella wangjianweii* (*IrdA and **IrdA, respectively). Purified reductases were assayed on a panel of identified electron acceptors. Ten purified reductases reduced at least one electron acceptor, with substrate specificity strongly tracking with the observed induction patterns (**Fig. 4E**). To further validate assigned substrate specificities, we tested whether *irdA* predicted strain-level variability in *E. lenta* itaconate utilization. Indeed, we found that *irdA* presence/absence predicted itaconate reduction across 5 different *E. lenta* strains (**Extended Data Fig. 10**).

Substrate-profiling experiments revealed particularly marked distinctions in cinnamate reductase substrate specificity. While the ability of *E. lenta* and *H. filiformis* to utilize diverse cinnamates could plausibly reflect the activity of a single promiscuous reductase, we found that both microbes encode multiple reductases with distinct specificities for different subsets of cinnamates (**Fig. 4E**). Among *H. filiformis* reductases, CirA utilized *p*-coumarate, chlorogenate, and rosmarinate. CirC weakly utilized cinnamate, while CirD utilized cinnamate, *m*-coumarate, *p*-coumarate, ferulate, and caffeate.

Intriguingly, while the cinnamate reductase CirB utilized *p*-coumarate, caffeate, chlorogenate, and rosmarinate, it also utilized the alkene resveratrol. Among *E. lenta* reductases, CrdC and CrdD both utilized sinapate, ferulate and caffeate, with CrdC uniquely using *p*-coumarate. These results demonstrate that *E. lenta* and *H. filiformis* express multiple cinnamate reductases with variable substrate specificities, some of which discriminate between substrates with relatively subtle structural differences.

### Reductases exhibit evolutionary complexity

We next explored the relationship between reductase evolution and substrate specificity. We reasoned that new reductase activities could be acquired either by (1) horizontal gene transfer or (2) gene duplication followed by functional divergence, and that these two scenarios would lead to distinct relationships between reductase sequence and substrate specificity. To investigate this, we constructed a phylogenetic tree using flavin reductases from *E. lenta*, *S. wadsworthensis*, and *H. filiformis* genomes (**Fig. 5**).

**Figure 5.**
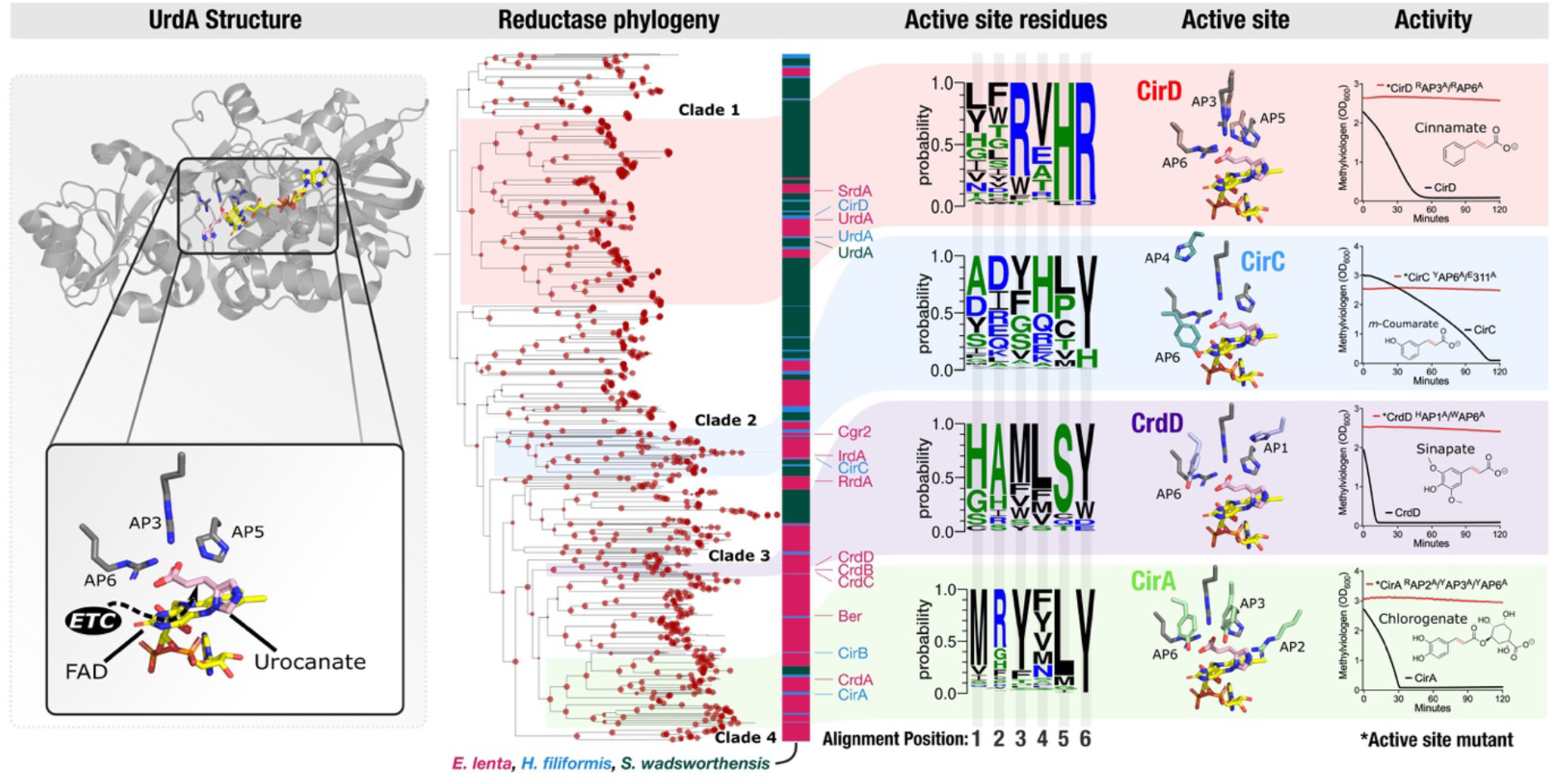
Independent evolutionary trajectories and distinct active sites distinguish flavin reductases with related electron acceptors. UrdA structure – previously published crystal structure of the *S. oneidensis* UrdA in complex with urocanate and flavin adenine dinucleotide (PDB code 6T87). Reductase phylogeny – phylogenetic tree of flavin reductases from *E. lenta*, *S. wadsworthensis*, and *H. filiformis* genomes. Bootstrap support values are indicated by the size of red dots at nodes of the tree and range from 70 to 100. Active site residues – representation of sequence identity of active site amino acids in reductase clades 1-4 scaled to frequency within the multiple sequence alignment. Positions within the multiple sequence alignment have been renumbered to active site alignment position (AP). Active site – AlphaFold models of CirD, CirC, CrdD, and CirA cinnamate reductases superimposed to the UrdA crystal structure. Activity –reductase activity of CirD, CirC, CrdD, and CirA and active site point mutants (indicated by an asterisk). Active site mutations and alignment positions they correspond to: *CrdD H313A (AP1) and W510A (AP6); *CirA R417A (AP2), Y469A (AP3), and Y634A (AP6); *CirC E311A and Y511A (AP6); *CirD R542A (AP3) and R716A (AP6). The y-axis shows the amount of reduced methylviologen in the presence of the indicated electron acceptor.

Several branches in the resulting tree include reductases from multiple species. For example, urocanate reductases from the three species are monophyletic and thus presumably share a more recent common evolutionary history (**Fig. 5**). However, a majority of branches on the tree are highly sub-branched and exclusively contain reductases from a single genus. These patterns suggest that both horizontal gene transfer and gene duplication may have played significant roles in reductase evolution within high reductase per genome bacterial lineages.

The distribution of reductases with different activities in the tree suggests a convoluted evolutionary history, with cinnamate reductases providing a striking example of the complex relationship between amino acid sequence and substrate specificity. Despite catalyzing highly similar reactions, we observed that the eight *E. lenta* and *H. filiformis* reductases induced in the presence of different cinnamate substrates separated into four phylogenetically distinct reductase clades (**Fig. 5, clades 1-4**). Cinnamate reductases typically share >30% amino acid sequence identity to other reductases within their clade – including reductases with distinct substrate specificities (*e.g.*, *H. filiformis* cinnamate reductase CirD and urocanate reductase UrdA from ‘reductase clade 1’ share 32% sequence identity). By contrast, cinnamate reductases from different reductase clades share <26% sequence identity, despite utilizing similar substrates (**Fig. 5**). These observations suggest that flavin reductases with similar cinnamate substrate specificities independently evolved at least four times.

### Dissimilar reductase active sites catalyze related reactions

To clarify how different evolutionary trajectories may have independently generated reductases with similar cinnamate substrate specificities, we turned to previous mechanistic studies of enzymes from the flavin reductase superfamily. Fumarate and urocanate reductases were previously shown to contain two conserved active site arginines – one that forms a critical salt bridge with the substrate carboxyl group and a second proposed to facilitate catalytic proton transfer to reduce the substrate enoate group.^37–39^ As a similar substrate enoate group is reduced by cinnamate reductases, we compared the substrate-bound urocanate reductase crystal structure to AlphaFold structural models of cinnamate reductases from the four reductase clades (**Fig. 5**). We found that active site arginines were conserved in reductase clade 1, including in the UrdA urocanate reductases and the CirD cinnamate reductase (**Fig. 5**, alignment positions 4 and 6). By contrast, arginines were not conserved at the same positions in reductase clades 2-4. Instead, tyrosine was conserved at alignment position 6 and other active site amino acids exhibited variable, clade-specific patterns of conservation (**Fig. 5**).

To test whether the distinct patterns of active site conservation in the four reductase clades might reflect mechanistic distinctions, we generated point mutants that targeted conserved amino acids in representative cinnamate reductases (CirA, CirC, CirD, and CrdD) from reductase clades 1-4. In each case, we found that conserved clade-specific active site amino acids were essential for activity (**Fig. 5**). These results thus show that distinct active site architectures functionally distinguish the cinnamate reductase clades and suggest that parallel evolutionary processes produced flavin reductases with similar substrate specificities but significant functional distinctions (see **Supplementary Discussion** for additional analysis).

## Discussion

Here we identify taxonomically distinct members of the gut microbiome that possess high numbers of respiratory-like reductases per genome. We find that these bacteria possess respiratory metabolisms and identify organic respiratory electron acceptors that exhibit strain-dependent patterns of usage. While much remains to be explored about the role of these activities, these results suggest a distinct type of respiration, defined by the versatile use of diverse organic respiratory electron acceptors, may play important roles within the gut.

Here we identify taxonomically distinct members of the gut microbiome that possess high numbers of respiratory-like reductases per genome that enable respiratory growth. Notably, these bacteria possess reductase arsenals (often >50 reductase/genome) that stand apart from previously characterized (non-host associated) bacterial respiratory specialists. For example, the marine and fresh-water-inhabiting bacterium *Shewanella oneidensis* is widely cited as a model organism with exceptionally broad respiratory capabilities, but “only” possesses 22 molybdopterin and flavin reductases (**Supplementary Table 2**).^40–42^

Perhaps related to this observation, most characterized respiratory electron acceptors used by non-host associated respiratory specialists like *S. oneidensis* are small inorganic compounds. By contrast, the respiratory electron acceptors identified in our studies are primarily organic metabolites. Host feeding produces a consistent influx of organic compounds that likely distinguishes the gut from many other microbial ecosystems. It thus stands to reason that the dramatic reductase expansion in gut microbes may reflect the enzymatic diversity required to match the high concentration of chemically complex metabolite electron acceptors present in the gut. These observations suggest that ecosystem-dependent ecological differences may explain the emergence of distinct respiratory strategies defined by (1) a large reductase arsenal and respiration of diverse metabolite electron acceptors versus (2) a smaller reductase arsenal and respiration of limited inorganic compounds.

A particularly important aspect of the identified respiratory metabolisms identified in our studies concerns their implication for the gut metabolome. Microbiome-derived imidazole propionate, the product of urocanate reduction, impairs insulin signaling through mTORC1 and is elevated in the serum of type 2 diabetes patients.^43^ Microbiome-derived hydro-*p*-coumarate (also called 4-hydroxyphenylpropionate or desaminotyrosine) protects against influenza through type I interferon activation.^44,45^ Resveratrol impacts multiple enzymes and transcription factors and influences several metabolic and immune pathways.^33^

### Further pointing to the significance of respiration in the metabolism of bioactive metabolites, several

*E. lenta* reductases with electron acceptors relevant for mammalian physiology have been previously described. Cgr2 is a flavin reductase that reduces the cardiac medication digoxin and an unidentified molecule that drives intestinal inflammation through Th17 activation.^36,46–48^ The flavin reductase Ber catalyzes a carbon-oxygen bond cleavage as part of a pathway that generates enterodiol and enterolactone, compounds that function similar to the endogenous hormone oestrogen.^49^ Finally, Dadh, is a molybdopterin reductase that dehydroxylates the neurotransmitter dopamine and inactivates the Parkinson’s disease medication L-DOPA.^9,50^

Itaconate is perhaps the most intriguing electron acceptor identified in our studies. In macrophages, itaconate is produced by the interferon-inducible gene Irg1 in response to bacterial and viral infection. In this context, itaconate acts as an immunometabolite that orchestrates direct and indirect mechanisms of pathogen defense.^33–35^ Several intracellular bacterial pathogens counteract mammalian itaconate by degrading it to pyruvate and acetyl-CoA.^51,52^ The reduction of itaconate to 2-methylsuccinate by itaconate reductase represents a second type of bacterial transformation of itaconate. 2-methylsuccinate has received little previous research attention, though a recent preprint reported that levels of 2-methylsuccinate and itaconate increase in the bile following infection with multiple intestinal bacterial pathogens.^53^ While, to our knowledge, neither itaconate nor 2-methylsuccinate have been studied within the gut, it is striking that both compounds are dramatically reduced in fecal samples from an antibiotic-treated population (**Fig. 3D**).

In conclusion, our studies establish a respiratory strategy defined by the use of a large reductase arsenal to access diverse metabolite electron acceptors. We show that reactions catalyzed by respiratory reductases effect various bioactive metabolites. Considering that the vast majority of reductases encoded by gut bacteria remain functionally uncharacterized, identified metabolisms may only scratch the surface of interactions between respiratory reductases and the gut metabolome. Continued study of respiratory electron acceptor usage may thus provide and important avenue for informing our understanding of the functional capacity and metabolic output of the gut microbiome.

## Methods

### Detection of molybdopterin and flavin reductases

Genomes and metagenome-assembled genomes of common prokaryotes in the human gut microbiome were downloaded from the version 2.0 of the Unified Human Gastrointestinal Genome (UHGG) in MGnify database on July 2022.^54^ Only species representative genomes identified in more than five distinct samples (1533 total) were included in subsequent analyses. Gene prediction was done with Prodigal (version 2.6.3)^55^ in single mode. Molybdopterin and flavin reductases were identified by searching PFAM Hidden Markov models (PF00384 and PF00890, respectively) with the tool Hmmsearch (version 3.3) (E-value cut-off 0.001).^56,57^ SIGNALP (version 5.0) was run to predict the putative cellular localization of the proteins using the parameter -org gram.^58^

To extend the genomic reductase content analyses to the identified taxonomic families, selected representative genomes for Burkholderiaceae, Eggerthellaceae and Erysipelotrichaceae (**Extended Data Fig. 1A-C, Supplementary Table 3**) were downloaded from the Genome Taxonomy Database on October 2022 (release 202 of April 27, 2021). The same protocol as described above was performed to detect the flavin and molybdopterin reductases.

### Concatenated ribosomal proteins phylogeny

Maximum-likelihood trees were calculated based on the concatenation of 14 ribosomal proteins (L2, L3, L4, L5, L6, L14, L15, L18, L22, L24, S3, S8, S17, and S19) for the UHGG tree and 16 ribosomal proteins for the three family trees (L16 and S10). Homologous protein sequences were aligned using MAFFT (version 7.467) (--auto option), and alignments refined to remove gapped regions using Trimal (version 1.4.1) (--gappyout option).^59,60^ The protein alignments were concatenated and manually inspected. Phylogenetic trees were inferred using using IQ-TREE with 1000 ultrafast bootstrap replicates (-bnni -m TEST -st AA -bb 1000 -nt AUTO) (version 1.6.12), using ModelFinder to select the best model of evolution, and with 1000 ultrafast bootstrap.^61–63^

### Reductase pangenomics

Pangenomes for *E*. *lenta*, *S*. *wadsworthensis*, and *H*. *filiformis* were calculated anvi’o 7.1 (with parameters anvi-pan-genome --mcl-inflation 10).^64^ Briefly, the program (1) performed an all vs. all NCBI BLAST to create a sequence similarity network, (2) used the Markov Cluster algorithm to resolve gene clusters, and (3) was visualized with “anvi-display-pan”.^65^ Subsequently, the species pangenomes were subsetted for gene clusters annotated with the Pfam PF00890 using “anvi-split”.

### General bacterial culture conditions

All bacterial strains used in this study are listed in **Supplementary Table 11**. Anaerobic growth was performed in a Coy vinyl anaerobic chamber under an atmosphere of 2-5% H_2_, 2-5% CO_2_, and balance nitrogen. Strains used in this study were initially grown on BHI media (BD BACTO Brain Heart Infusion #237500) plates for 48-72 hours at 37 °C to yield visible colonies. Overnight cultures were grown for 24 hours at 37 °C using liquid BHI for *E. lenta* and *S. wadsworthensis*, or liquid BHI-NoDex (Alpha Bioscience Brain Heart Infusion W/O Dextrose #B02-114) plus 0.01% tween-80 for *H. filiformis*.

### ATP assay

Bacterial cultures were grown and conditioned overnight in 10 mM sodium formate and 10 mM electron acceptor to induce reductase expression. Cells were washed and resuspended in phosphate assay buffer (40 mM potassium phosphate, 10 mM magnesium sulfate, pH 7.0) then allowed to incubate at room temperature for 45 minutes to achieve a resting cell suspension. To initiate the reaction, cells were mixed 1:1 with buffer containing 2 mM sodium formate or buffer containing 2 mM sodium formate additionally supplemented with 2 mM electron acceptor in a 96-well plate. At specified time points 10 µL of mixture was taken out and placed into a separate 96-well plate with 90 µL DMSO to quench the reaction and liberate cellular ATP. The ATP concentration was measured by using an ATP determination kit (Invitrogen #A22066). Sample luminescence was read using black-well plates (Corning 96-well Black #3915) in a BioTek Cytation 5 plate reader and compared to a standard curve of known ATP concentrations.

### Respiratory electron acceptor screen

A collection of potential respiratory electron acceptors were selected for testing based on: 1) their likely presence in the gut, 2) commercial availability, and 3) previous reports of their reduction by species in the Eggerthellaceae family (in the case of some phenylpropanoid derivatives).^9,49,66^ Selected potential electron acceptors were directly dissolved into media at 10 mM in either 50% BHI media containing 10 mM sodium formate (*E. lenta* and *S. wadsworthensis*), or 100% BHI-NoDex with 0.01% tween-80 (*H. filiformis*) and filter sterilized. Molecules were not dissolved in solvents first to avoid potential issues of their additional usage by bacteria. Media was arrayed into deep-well (1 mL) 96-well plates (USA Scientific PlateOne #1896-1110) for growth. Strains were inoculated with normalized 1/1000 inoculations from overnight cultures, and plates were sealed with adhesive seals to minimize evaporation. Time points were taken at 24, 48, 72, and 96 hours by mixing with a multichannel pipette and moving 200 µl into a separate 96-well plate (Greiner Bio-One 96-Well Microplates #658162) and reading absorbance at OD_600_ in a BioTek Cytation 5 plate reader normalized to media blanks.

### Metabolite extraction from liquid culture

Samples were incubated at −80 °C for between one and twelve hours. Four volumes of methanol containing internal standards (d^7,15^N-proline, Cambridge Isotopes, DNLM-7562 and U-^13^C-palmitic acid, Cambridge Isotopes, CLM-409) were added to each culture supernatant (one volume) in a microcentrifuge tube. Tubes were then centrifuged at −10 °C, 20,000 x *g* for 15 min followed by transfer of 100 μL of supernatant to pre-labeled mass spectrometer autosampler vials (Microliter, 09-1200).

### Metabolite extraction from fecal material

Metabolites were extracted from fecal samples by adding 80% methanol (containing ^13^C_9_-phenylalanine, Cambridge Isotopes, CLM-2250; d_6_-succinate, DLM-831; and ^13^C_11_-tryptophan CLM-4290 internal standards) to 100 mg/mL and stored at −80 °C for at least one hour in beadruptor tubes (Fisherbrand; 15-340-154). Samples were then homogenized at 4 °C on a Bead Mill 24 Homogenizer (Fisher; 15-340-163) set at 1.6 m/s with six thirty-second cycles, five seconds off per cycle. Samples were then centrifuged at −10 °C, 20,000 x *g* for 15 min to generate supernatants for subsequent metabolomic analysis.

### Derivatization methods for GC-MS analyses

Culture supernatants were dried completely under nitrogen stream at 30 °C (Biotage SPE Dry 96 Dual, 3579M). To dried samples, 50 μL of freshly prepared 20 mg/mL methoxamine (Sigma, 226904) in pyridine (Sigma, 270970) was added and incubated in a ThermoMixer (Eppendorf) for 90 min at 30 °C and 1400 rpm. After samples were cooled to room temperature, 80 µL of derivatizing reagent (N,O-Bis(trimethylsilyl)trifluoroacetamide, 1% Trimethylchlorosilane, Sigma, B-023) and 70 μL of ethyl acetate (Sigma, 439169) were added and mixed in a ThermoMixer at 70 °C for one hour at 1400 rpm. Samples were cooled to room temperature and diluted with 400 μL of ethyl acetate for analysis.

### GC-MS chromatography and instrument parameters

Culture supernatants were analyzed using a GC-MS (Agilent 7890A GC system, Agilent 5975C MS detector) with an electron impact ionization source, a HP-5MSUI column (30 m x 0.25 mm, 0.25 µm, Agilent Technologies 19091S-433UI), and 1 μL injection. Oven ramp parameters: 1 min hold at 60 °C, 16 °C per min up to 300 °C with a 7 min hold at 300 °C, for a total runtime of 23 minutes. Inlet temperature was 280 °C, transfer line was 300 °C, and source temperature was 230 °C. Ionization was achieved using a 70 eV electron beam. Ions were measured over a 50 to 600 m/z range.

### GC-MS data analyses

Culture supernatant data analyses were performed using MassHunter Quantitative Analysis software (version B.10, Agilent Technologies). Metabolite identifications were confirmed by matching to authentic standard spectra and retention time and spectra in the NIST Tandem Mass Spectral Library Version 2.3 (see **Supplementary Tables 12 & 13** for background and validation methods of each metabolite). Thus, all metabolites reported from mouse fecal pellet samples and human stool samples are identified at confidence Level 1.^67^ Normalized peak areas were calculated by dividing raw peak areas of targeted analytes by averaged raw peak areas of internal standards d7-proline and U-^13^C-palmitate.

### LC-MS parameters for culture supernatant analyses

Catechin, epicatechin, chlorogenate, rosmarinate, and shikimate reduction products were not readily available as standards were not readily available as standards, therefore LC-MS/MS fragmentation was used to identify putative products of these reduction reactions in microbial cultures. All LC-MS analyses were performed using a Thermo Scientific Vanquish Flex UHPLC coupled to an IQ-X mass spectrometer (Thermo Fisher Scientific). To detect these compounds and their putative reduced products in *E. lenta* and *S. wadsworthensis* cultures, the chromatographic method used was an isocratic 100% mobile phase A (5% acetonitrile, 0.1% formic acid in water) for 0.2 min, followed by a gradient of 0 to 97% mobile phase B (99.9% acetonitrile, 0.1% formic acid) for 4.5 min with a wash of 100% mobile phase B for 1 min. Catechin/epicatechin, chlorogenate, rosmarinate, and shikimate and their putative reduced products were detected by mass spectrometry using negative ionization. Flow from the UHPLC was ionized with the heated electrospray ionization (HESI) source set to

2400 V, ion transfer tube temp set to 200 °C, vaporizer temperature set to 400 °C, and sheath, aux and sweep gases set to arbitrary values of 40, 5, and 1, respectively. Data for MS^1^ was acquired using a maximum inject time of 50 ms, a normalized AGC target of 25%, a 100–1300 m/z quadrupole scan range, and a resolution of 60,000. All tandem MS^2^ mass spectral data was acquired using a 1.5 m/z quadrupole isolation window, a maximum inject time of 22 ms, a normalized AGC target of 20%, and a resolution of 15,000. Each of the metabolite ions of interest were added to an inclusion list for MS^2^ fragmentation by a normalized higher-energy collisional dissociation energy of 30%. Standards for catechin and epicatechin reduction products, hydrochlorogenate, hydrorosmarinate, and hydroshikimate (3,4,5-trihydroxycyclohexanecarboxylic acid)were not readily available, therefore, they are reported at confidence Level 3.^67^ To support the identification of the putative catechin and epicatechin reduction products, a comparison of the derivative MS^2^ fragmentation spectra to a representative catechin/epicatechin MS^2^ fragmentation spectrum has been included (**Supplementary Fig. 1 & 2**). To support the assignment of the hydrochlorogenate and hydrorosmarinate structures, analysis of the MS^2^ fragmentation spectrum and a comparison to the hydrochlorogenate and hydrorosmarinate MS^2^ fragmentation spectra, respectively, have been included with putative fragment ion structures (**Supplementary Fig. 3 & 4**. To support the assignment of the hydroshikimate structure, analysis of the MS^2^ fragmentation spectrum and a comparison to the shikimate MS^2^ fragmentation spectrum has been included with putative fragment ion structures (**Supplementary Fig. 5**).

### LC-MS parameters for fecal material analyses

All metabolites from mouse fecal pellet and human stool were measured by LC-MS analyses. Reversed-phase chromatography was performed at a 300 μl min−1 flow rate on a Waters CORTECS T3 C18 RP-UHPLC column (100 × 2.1 mm inner diameter, 1.6 μm particle size, 120 Å pore size (1 Å = 0.1 nm)). Mobile phase A was 5% acetonitrile in water with 0.1% formic acid and mobile phase B was acetonitrile with 0.1% formic acid. For all mouse fecal pellet and human stool analyses, the chromatographic method used was an isocratic 100% mobile phase A for 0.2 min, followed by a gradient of 0 to 40% mobile phase B for 2.8 min, then a gradient from 40% to 100% mobile phase B over 1.5 min, with a final wash of 100% mobile phase B for 1.5 min. To acquire negative ionization mass spectra, flow from the UHPLC was ionized with the heated electrospray ionization (HESI) source set to 3500 V, ion transfer tube temp set to 190 °C, vaporizer temperature set to 120 °C, and sheath, aux and sweep gases set to arbitrary values of 50, 10, and 1, respectively. For positive ionization mass spectra, flow from the UHPLC was ionized with the heated electrospray ionization (HESI) source set to 3500 V, ion transfer tube temp set to 300 °C, vaporizer temperature set to 350 °C, and sheath, aux and sweep gases set to arbitrary values of 50, 10, and 1, respectively. Data for both positive and negative mode MS and MS^2^ were acquired using the EASY-IC internal calibration to fluoranthene, automatic maximum inject time, and standard AGC target. Further, data for all MS^1^ was acquired using a 100–365 m/z quadrupole scan range and a resolution of 120,000. All tandem MS^2^ mass spectral data was acquired using a 0.7 m/z quadrupole isolation window and a resolution of 15,000. Each of the metabolite ions of interest were added to an inclusion list for MS^2^ fragmentation by stepped higher-energy collisional dissociation absolute energies of 20, 40, 60, and 80 eV.

### LC-MS data analysis

Mouse fecal pellet and human stool sample data analysis was performed using FreeStyle software (version 1.8 SP2, Thermo Scientific). Metabolite identification was established by matching accurate mass, retention time, and MS^2^ fragmentation pattern from candidate ions in experimental samples to data from authentic standards of each metabolite of interest. Thus, all metabolites reported from mouse fecal pellet samples and human stool samples are identified at confidence Level 1.^67^ Normalized peak areas were calculated by dividing raw peak areas of targeted analytes by averaged raw peak areas of internal standards (^13^C_9_-phenylalanine and ^13^C_11_-tryptophan for metabolites measured by positive ionization; d_6_-succinate and ^13^C_11_-tryptophan for metabolites measured by negative ionization).

### GC-MS and LC-MS Quality Control

Biological control plasma samples were processed and analyzed alongside culture supernatant samples to evaluate metabolite extraction efficiency and instrument performance. The samples were extracted with methanol containing known internal standard (IS) concentrations. Recovery, retention time, and coefficient of variation (CV) were calculated for ISs. Method blanks containing no metabolites were included throughout the run.

### Collection of mouse fecal samples

All mouse experiments were performed in accordance with and approved by the Institutional Animal Care and Use Committee of the University of Chicago under protocol 72599. Six- to nine-week-old female specific pathogen-free C57BL/6 mice from Jackson Laboratories were used for all experiments. Mice were kept within a facility that maintained a 12/12-hour light/dark cycle and controlled humidity (30-70%) and temperature (68-79 °F). Mice were housed in sterile, autoclaved cages with irradiated food and acidified, autoclaved water upon arriving at on site mouse facility. Mouse handling and cage changes were performed by investigators wearing sterile gowns, masks, and gloves in a sterile biosafety hood. Mice were cohoused with their original shipment group until after antibiotic treatment. Fresh fecal pellets were collected directly from mice.

Following collection of “pre-antibiotic” treatment fecal pellets, mice were administered metronidazole, neomycin, and vancomycin (MNV) each at 0.25 g/L in drinking water for 3 days. The drinking water was then changed to acidified, autoclaved water and the mice were transferred to clean cages. Mice were injected intraperitoneally with clindamycin 0.4 mg total, 2 days after the end of the MNV regimen and received a cage change each day following to prevent additional antibiotic consumption through coprophagy. “Post-antibiotic” treatment fecal pellets were collected 2 days after the clindamycin injection.

### Collection of human fecal samples

Fecal samples from healthy, non-antibiotic treated subjects were collected through a prospective fecal collection protocol (UC Protocol IRB20-1384) approved by the institutional review boards (IRB) at the University of Chicago (UC). All donors provided written and informed consent for IRB-approved biospecimen collection and analysis. The study was conducted in accordance with the Declaration of Helsinki.

Fecal samples from patients treated with broad-spectrum antibiotics were selected from samples collected as part of a prospective cohort study of hospitalized adult hepatology patients at a single institution.^68^ Inclusion criteria for the study included: age ≥18 years, ability to provide informed consent (either themselves or by proxy if decisionally incapacitated) and being treated on the hepatology consult service. Subjects who were younger than 18 years, unable to provide consent, had prior solid organ transplant, or a prior colectomy were excluded. Patients were enrolled as soon as possible upon hospital admission, most within 48 hours. All samples were obtained under a protocol that was approved by the University of Chicago IRB (IRB21-0327), and written informed consent was obtained from all participants or their surrogate decision makers.

Upon collection, fecal samples were refrigerated at +4°C for less than 24 hours prior to aliquoting and storing at −80°C until processing for shotgun metagenomic and metabolomic analysis (details below). Demographic and clinical data, including medication administration, were collected through review of the medical record and through the University of Chicago Center for Research Informatics (CRI).

The 22 samples from antibiotic-treated patients included in this study were chosen based on antibiotic administration and metagenomic findings that would be expected after broad-spectrum antibiotic exposure. The most common antibiotic exposure was a combination of intravenous (IV) vancomycin, IV cefepime, and IV metronidazole; however, one patient was also exposed to meropenem, two patients exposed to piperacillin-tazobactam, and one patient exposed to ciprofloxacin and ceftriaxone. The metagenomic changes included 8 samples with >90% relative abundance of Proteobacteria and 14 samples with >90% relative abundance of *Enterococcus*, which indicates that broad spectrum antibiotic exposure had marked effects on the gut microbiome composition in these samples.

### Fecal DNA extraction

DNA was extracted using the QIAamp PowerFecal Pro DNA kit (Qiagen). Prior to extraction, samples were subjected to mechanical disruption using a bead beating method. Briefly, samples were suspended in a bead tube (Qiagen) along with lysis buffer and loaded on a bead mill homogenizer (Fisherbrand). Samples were then centrifuged, and supernatant was resuspended in a reagent that effectively removed inhibitors. DNA was then purified routinely using a spin column filter membrane and quantified using Qubit.

### Metagenomic analyses of human fecal samples

Human fecal samples underwent shot-gun DNA sequencing. After undergoing mechanical disruptions with a bead beater (BioSpec Product), samples were further purified with QIAamp mini spin columns (Qiagen). Purified DNA was quantified with a Qubit 2.0 fluorometer and sequenced on the Illumina HiSeq platform, producing around 7 to 8 million PE reads per sample with read length of 150 bp. Adapters were trimmed off from the raw reads, and their quality was assessed and controlled using Trimmomatic (v.0.39), then human genome was removed by kneaddata (v0.7.10, https://github.com/biobakery/kneaddata).^69^ Taxonomy was profiled using metaphlan4.^70^ Alpha-diversity (a reflection of the number of unique bacterial taxa and their relative abundances) of fecal samples was estimated using Inverse Simpson Index. Raw sequencing data generated were deposited to the National Center for Biotechnology Information (NCBI) Sequence Read Archive (SRA) under accession number PRJNA912122.

### 16S amplicon sequencing analyses of mouse fecal samples

The V4-V5 region within 16S rRNA gene was amplified using universal bacterial primers – 563F (5’-nnnnnnnn-NNNNNNNNNNNN-AYTGGGYDTAAA-GNG-3’) and 926R (5’-nnnnnnnn-NNNNNNNNNNNN-CCGTCAATTYHT-TTRAGT-3’), where ‘N’ represents the barcodes, ‘n’ are additional nucleotides added to offset primer sequencing. Amplicons were then purified using magnetic beads, then quantified, and pooled at equimolar concentrations. Illumina sequencing-compatible Combinatorial Dual Index (CDI) adapters were ligated onto pooled amplicons using the QIAseq 1-step amplicon library kit (Qiagen). Library QC was performed using Qubit and Tapestation and sequenced on Illumina MiSeq platform to generate 2×250bp reads, generating 5000-10000 reads per sample. Raw V4-V5 16S rRNA gene sequence data is demultiplexed and processed through the dada2 pipeline into Amplicon Sequence Variants (ASVs) with minor modifications in R (v4.0.3). Specifically, reads were first trimmed at 210 bp for forward reads and 150 for reverse reads to remove low quality nucleotides. Chimeras were detected and removed using the default consensus method in the dada2 pipeline. Then, ASVs with length between 300 bp and 360 bp were kept and deemed as high quality ASVs. Taxonomy of the resultant ASVs were assigned to the genus level using the RDP classifier (v2.13) with a minimum bootstrap confidence score of 80. Species-level classification can be provided using blastn (v2.13.0) and refseq_rna database (updated 2022-06-10).

### Transcriptomics analyses

*E. lenta* and *H. filiformis* were grown in 50 mL of media supplemented with 10 mM formate and 10 mM of the tested electron acceptor. A BHI baseline media was used for *E. lenta* cultures. A no-dextrose BHI media supplemented with 0.01% tween-80 was utilized for *H. filiformis* cultures. Overnight cultures were diluted 1/100 into flasks and allowed to grow shaking for 24 hours. Whole cultures were pelleted and used for RNA extraction. Total RNA from biological replicates was extracted using the Maxwell RSC instrument (Promega). RNA was quantified using Qubit, a fluorometric method and integrity was measured through a TapeStation, Agilent Technologies. Ribosomal RNA was removed using NEBNext rRNA Depletion Kit for bacteria. Libraries from ribo-depleted samples were constructed using the NEB Ultra Directional RNA library prep kit for Illumina. Briefly, 10-500ng total RNA was subjected to ribosomal RNA depletion and rRNA-free samples were fragmented based on their RNA integrity number. Post cDNA synthesis, Illumina compatible adapters were ligated onto the inserts and final library quality was evaluated using TapeStation (Agilent technologies). Libraries were normalized using qPCR and then sequenced on Illumina’s NovaSeq 6000 platform using 2×150bp read configuration.

High quality sequence reads were mapped to the NCBI reference genomes of *E. lenta* strain DSM2243 (GCF_000024265.1), and *H. filliformis* strain AF24-29 (GCF_003459085.1), accordingly. The reads were mapped to the respective reference genomes using Bowtie2 v2.4.5 and sorted with Samtools v1.6 using default parameters.^71,72^ All sequence data were deposited to the NCBI SRA database under the Bioproject ID PRJNA1012298. Read counts were generated using featureCounts v2.0.1 with the flags -p, -B, and -C.^73^ Gene expression was quantified as the total number of reads uniquely aligning to the respective reference genomes, binned by annotated gene coordinates.

Differential gene expression and related quality control analyses were determined using the Bioconductor package DESEQ2 in R.^74,75^ Normalization of raw read counts was performed by a scaling method implemented within DESEQ2 package, which accounts for differences in library size and library composition. Reads were also normalized for batch effects where applicable by ComBat-seq.^76^ Log_2_-fold change shrinkage was performed using apgelm method in DESeq2.^77^ Differential expression of pairwise comparisons (of the different conditions) was assessed using the negative binomial test with a Benjamini–Hochberg false discovery rate (FDR) adjustment applied for multiple testing corrections.

### Proteomics mass spectrometry

Samples for proteomic analysis were prepped using a modified version of the filter-aided sample preparation (FASP) procedure. A portion of thawed cell pellet was mixed with 500 µL 1x SDS buffer (40 mL of 1x contained: 4g glycerol, 0.67g Tris HCl, 0.68g Tris base, 0.8g SDS, and 6mg EDTA) and dithiothreitol to 20 mM. Mixture was sonicated at 20% pulsing 2-sec on 2-sec off for 5 minutes. This mixture was then heated at 95 °C for 20 minutes, followed by 37 °C for 30 minutes. Iodoacetamide was added to 60 mM and mixture was incubated in the dark for 60 minutes, before dithiothreitol was again added to 60 mM. This prepared lysate was then mixed 1:8 (25µl:200µl) with exchange buffer (8 M urea, 0.2% (w/v) deoxycholate, 1M ammonium bicarbonate pH 8) and dispensed to a filter unit. Mixture was spun at 14,000g for 10 minutes, discarding filtrate. The filter unit was washed 3 times each with 200 µL exchange buffer and spun at 14,000g for 10 minutes discarding the filtrate. The filter unit was then washed twice with 200 µL digestion buffer (0.2% (w/v) deoxycholate, 50 mM ammonium bicarbonate pH 8) and spun at 14,000g for 10 minutes discarding the filtrate. The filter unit was transferred to a passive collection tube and 100 µL digestion buffer + trypsin (∼1:50 trypsin:protein) and incubated at 37 °C overnight. The following day, the tube was centrifuged at 14,000g for 10 minutes, but filtrate was not discarded. 50 µL of peptide recovery buffer (50 mM ammonium bicarbonate pH 8) was added to the filter unit and spun at 14,000g for 10 minutes, and repeated once more. The resulting three filtrates were transfer to a LoBind tube. To the filtrate, 900 µL of ethyl acetate and 2.5 µL of trifluoroacetic acid were added and vortexed. Mixture was sonicated (10 seconds at 10%), and centrifuged at 16,000g for 10 minutes. The upper organic layer was removed and discarded, taking care not to disturb the phase boundary. The addition of ethyl acetate, sonication, centrifugation, and layer removal was repeated twice more, but without the addition of trifluoroacetic acid. Sample tubes were heated uncovered to 60 °C for 5 minutes to evaporate residual ethyl acetate, then frozen at −80 °C until analysis.

Samples of trypsin-digested proteins were analyzed using a Synapt G2-Si ion mobility mass spectrometer that was equipped with a nanoelectrospray ionization source (Waters, Milford, MA). The mass spectrometer was connected in line with an Acquity M-class ultra-performance liquid chromatography system that was equipped with trapping (Symmetry C18, inner diameter: 180 μm, length: 20 mm, particle size: 5 μm) and analytical (HSS T3, inner diameter: 75 μm, length: 250 mm, particle size: 1.8 μm) columns (Waters). The mobile phase solvents were water and acetonitrile, both of which contained 0.1% (volume/volume) formic acid. Data-independent, ion mobility-enabled, high-definition mass spectra and tandem mass spectra were acquired in the positive ion mode.^78–81^

Data acquisition was controlled using MassLynx software (version 4.1) and tryptic peptide identification and relative quantification using a label-free approach were performed using Progenesis QI for Proteomics software (version 4.0, Waters).^82^ Data were searched against the *E*. *lenta* protein database to identify tryptic peptides.^83^

### Recombinant reductase production and purification

Hexa-histidine-tagged reductase constructs were expressed in *E. coli* Rosetta cells using either the pET28a or pMCSG53 expression vectors, as previously described.^84^ Briefly, expression was performed using 500 mL cultures of Luria-Bertani broth (BD LB Broth, Miller #244610) + 100 µM riboflavin, grown to an OD of 0.7-1.0 at 37 °C, then induced with 1 mM β-d-1-thiogalactopyranoside overnight at 20 °C. Kanamycin (30 mg/mL, pET28a vector) or carbenicillin (100 mg/mL, pMCSG53 vector) were used for selection. Cultures were pelleted, resuspended in lysis buffer (50 mM Tris-HCl, 300 mM NaCl, 1 mM dithiothreitol, 10 mM imidazole, pH 7.5), and incubated with 500 µg/mL lysozyme on ice for 30 minutes. Cells were ruptured by sonication (Branson Digital Sonifier SFX 250) with 40 15-second pulses between 40-second waits. Insoluble material was removed by centrifugation. Resulting supernatants were mixed with Ni-charged resin beads (Bio-Rad Profinity IMAC) to purify the His-tagged reductases. Beads were washed using lysis buffer, and bound protein was eluted using lysis buffer containing an additional 500 mM imidazole. The eluted protein was dialyzed (Slide-A-Lyzer MINI Dialysis Device, 7K MWCO) into 50 mM HEPES, 100 mM NaCl, pH 7.0 for subsequent assays.

### Recombinant reductase assay

Assays were performed in an anaerobic chamber under an atmosphere of 2-5% H_2_, 2-5% CO_2_, and balance nitrogen. Potential electron acceptors were dried in 96-well plates (Falcon 96-well Polystyrene Microplates #351172) to allow for a 1 mM final concentration at a 200 µL reaction volume. Reaction buffer consisted of a base buffer of 50 mM HEPES and 100 mM NaCl pH 7.0, was autoclaved then immediately sparged with nitrogen and placed into the anaerobic chamber. Powders of methyl viologen (CAS #75365-73-0), sodium dithionite (CAS #7775-14-6), and flavin adenine dinucleotide (CAS #146-14-5) were weighted out and placed into the anaerobic chamber 24 hours before use. All components (plastics, buffers, etc.) were placed in the anaerobic chamber for at least 24 hours prior to assay use. Powders were resuspended in the base buffer immediately before usage. Methyl viologen was resuspended to 50 mM, sodium dithionite to 25 mM, and flavin adenine dinucleotide to 1 mM. The working master mix consisted of both methyl viologen and sodium dithionite added to 100 µM, which yields a deep purple color. flavin adenine dinucleotide was added to 40 µM, the deep purple color should remain. Purified reductases were added to a final concentration of 5 µg/well, and the resulting mixture was multichannel pipetted into the electron acceptor containing plates. Assays of CirA, CirC, CirD, CrdD and their point mutants used reductases at 0.5, 15, 5, 0.5 µg/well, respectively. Resuspension of the dried electron acceptor molecules begins the reaction. The plates were adhesive-sealed and read in a BioTek Epoch 2 at OD_600_ for 2 hours, where a reduction in absorbance indicates a consumption of methyl viologen as an electron donor in the reaction.

### Reductase phylogenetics

Genomes for *E. lenta*, *S. wadworthensis*, and *H. filiformis* were downloaded using ncbi-genome-download (https://github.com/kblin/ncbi-genome-download) (see **Supplementary Table 14** for genome accessions). We then used anvi’o v7.1 to convert genome FASTA files into contigs databases (https://anvio.org/m/contigs-db) using the contigs workflow (https://doi.org/10.1186/s13059-020-02195-w), during which Prodigal v2.6.3 identified open reading frames (contigs workflow).^55,85^ Next, we used the EcoPhylo workflow implemented in anvi’o (https://anvio.org/m/ecophylo) in ‘tree-mode’ to recover reductase genes in genomes using the Pfam model PF00890 and to explain their phylogeny. Briefly, the EcoPhylo workflow (1) used the program hmmsearch in HMMER v3.3.2^86^ to identify reductases, (2) removed hits that had less than 80% model coverage to minimize the likelihood of false positives due to partial hits, (3) dereplicated resulting sequences using MMseqs2 13.45111^87^ to avoid redundancy, (4) calculated a multiple sequence alignment with MUSCLE v3.8.1551,^88^ trimmed the alignment by removing columns of the alignment with trimal v1.4.rev15 (with the parameters ‘-gappyout’)^60^ (5) removed sequences that have more than 50% gaps using the anvi’o program anvi-script-reformat-fasta, (6) calculated a phylogenetic tree with the final alignment with IQ-TREE 2.2.0-beta COVID-edition^89^ (with the parameters ‘-nt AUTO -m WAG -B 1000’) that resulted in a NEWICK formatted tree file, and finally (7) visualized the tree in the anvi’o interactive interface.

### Reductase active site conservation analyses

AlphaFold models of CirA, CirC, CirD, and CrdD were downloaded from Uniprot.^90^ Active site amino acids were identified by independently superimposing N- and C-terminal domains of AlphaFold models to the substrate-bound urocanate reductase crystal structure (PDB code 6T87) using PyMOL v2.5.1 (http://www.pymol.org/pymol) ‘align’.^39^ Next, sequences in the monophyletic clade surrounding the experimentally validated sequences were subsetted from the reductase multiple sequence alignment using the program anvi-script-reformat-fasta and alignment positions were sliced in Jalview v2.11.2.5).^91^ Finally, the extent of conservancy among the active site-associated residues were visualized with WebLogo 3 (https://weblogo.threeplusone.com/create.cgi).

### Statistics and reproducibility

Statistical methods were not used to predetermine sample size. Due to impracticality of experiment blinding, experiments were not randomized and the investigators were not blinded. However, quantitative metrics were used to measure all results reducing the risk of bias. Data distribution was assumed to be normal but this was not formally tested.

## Data Availability

The datasets generated within the current study are available within the paper and the supplementary information. Quantitative metabolomic raw data files will be publicly available at the time of publication on MassIVE repository (ID MSV000093291). Transcriptomic datasets generated in this study can be found under Bioproject PRJNA1012298. Metagenomic data used in this study are publicly available on NCBI under BioProject IDs PRJNA912122 and PRJNA838648. Proteomic datasets can be accessed at doi.org/10.7910/DVN/OY9MPE. NCBI accession codes for studied proteins are provided in Supplementary Table 10.

## Acknowledgments

We thank Huaiying Lin & Nicholas P. Dylla for assistance with data analyses and Laurie Comstock for helpful feedback. We thank the University of Chicago Animal Resources Center for their assistance with mouse work (RRID:SCR_021806). Research reported in this publication was supported by funding from the National Institutes of Health (T32DK007074 to M.A.O., 1S10OD020062-01 to A.T.I., and K22AI144031 & R35GM146969 to S.H.L) and the Searle Scholars Program (to S.H.L).

## Author contributions

A.S.L., E.G.P., and S.H.L. conceptualized the project. A.S.L., I.T.Y., P.N.B., J.S., K.S., and D.S. performed experiments. A.S.L. M.S.S., P.N.B., and S.H.L. analyzed data. A.S.L., I.T.Y., P.N.B., J.S., and K.S. performed growth assays. A.S.L. performed ATP determination assays. A.S.L. and J.S. performed protein expression and purification, and reductase activity assays. M.S.S., R.M., and A.M.E. performed bioinformatic analysis, including phylogeny and pangenomes. A.S.L., R.R., R.S., and A.S. performed bioinformatics analysis of transcriptomic data. M.W.M., W.L., D.M., M.M., A.M.S. performed and analyzed mass spectrometry. A.T.I. performed and analyzed proteomics data. A.S.L., P.N.B., J.S., and E.W. performed animal experiments and maintenance. M.A.O. provided human fecal samples for analysis. A.S.L. and S.H.L. wrote the manuscript.

---

**Extended Data Figure 1.**
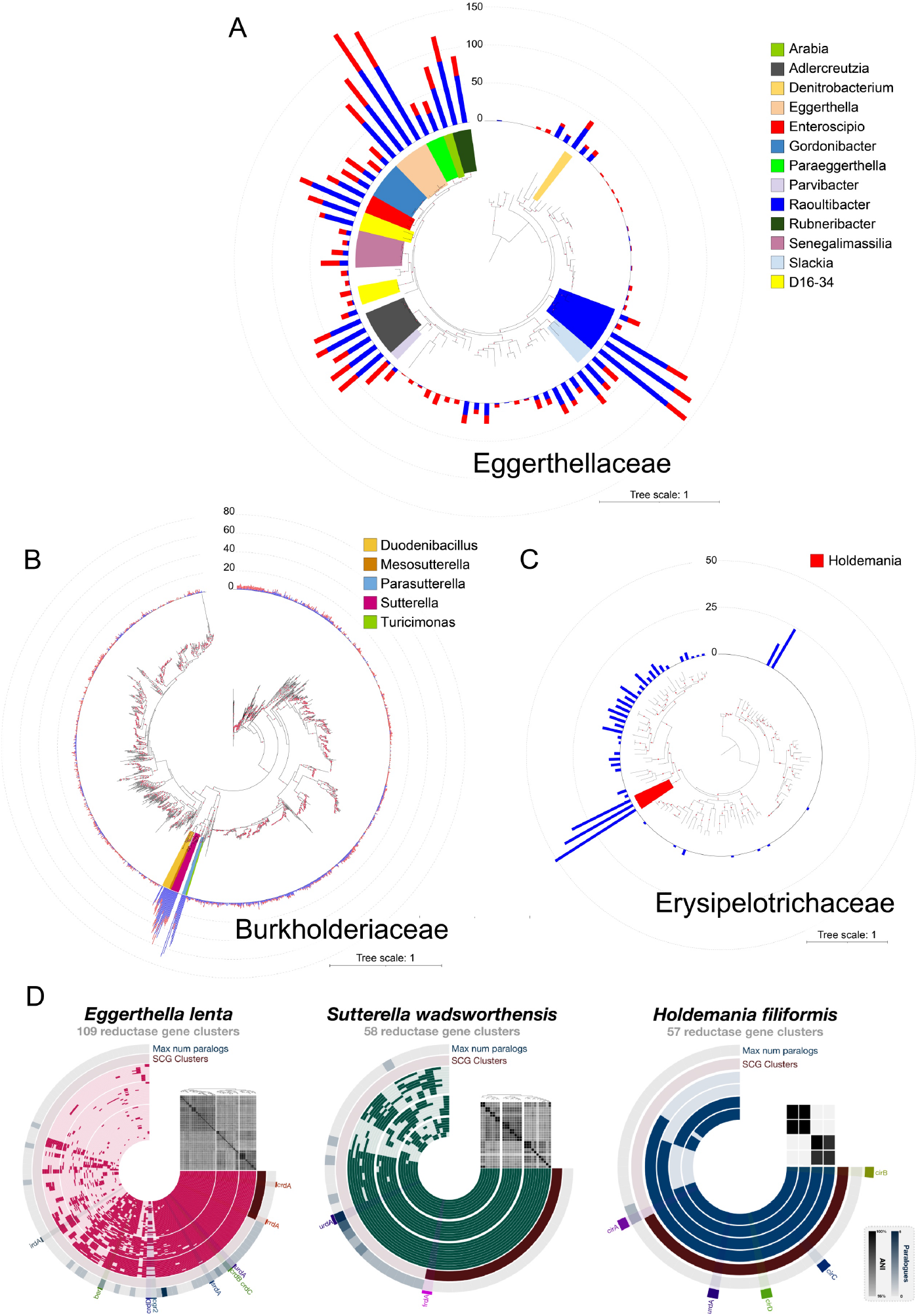
Distribution and identity of flavin and molybdopterin reductases in three taxonomic families. Phylogenetic trees constructed with representative genomes from each Genome Taxonomy Database (GTDB) species in (A) Eggerthellaceae (88 genomes, 2387 amino acid sites), (B) Burkholderiaceae (1510 genomes, 2379 amino acid sites), and (C) Erysipelotrichaceae (116 genomes, 2464 amino acid sites). Each maximum likelihood tree was constructed based on a concatenated alignment of 16 ribosomal proteins under an LG + I + G4 model of evolution. The numbers of flavin (blue) and molybdopterin (red) reductases with a computationally predicted signal peptide in each genome are graphed on the outer ring of the trees. (D) Flavin reductase pangenomes of *E. lenta*, *S. wadsworthensis*, and *H. filiformis*. For each pangenome, the inner concentric layers represent unique genomes while the radial elements represent gene cluster presence (darker color) or absence (lighter color) across the genomes. The outermost concentric circle, “Max num paralogues,” indicates the maximum number of paralogues (defined as reductases with ∼60% sequence identity) one genome contributes to the gene cluster. The second outermost circle, “SCG clusters,” indicates single-copy core reductase i.e., gene clusters for which every genome contributed exactly one gene. Genomes (inner concentric layers) are clustered by the presence/absence of reductase gene clusters. All vs all genome average nucleotide identity is depicted in the heat map above the genome concentric layers.

**Extended Data Figure 2.**
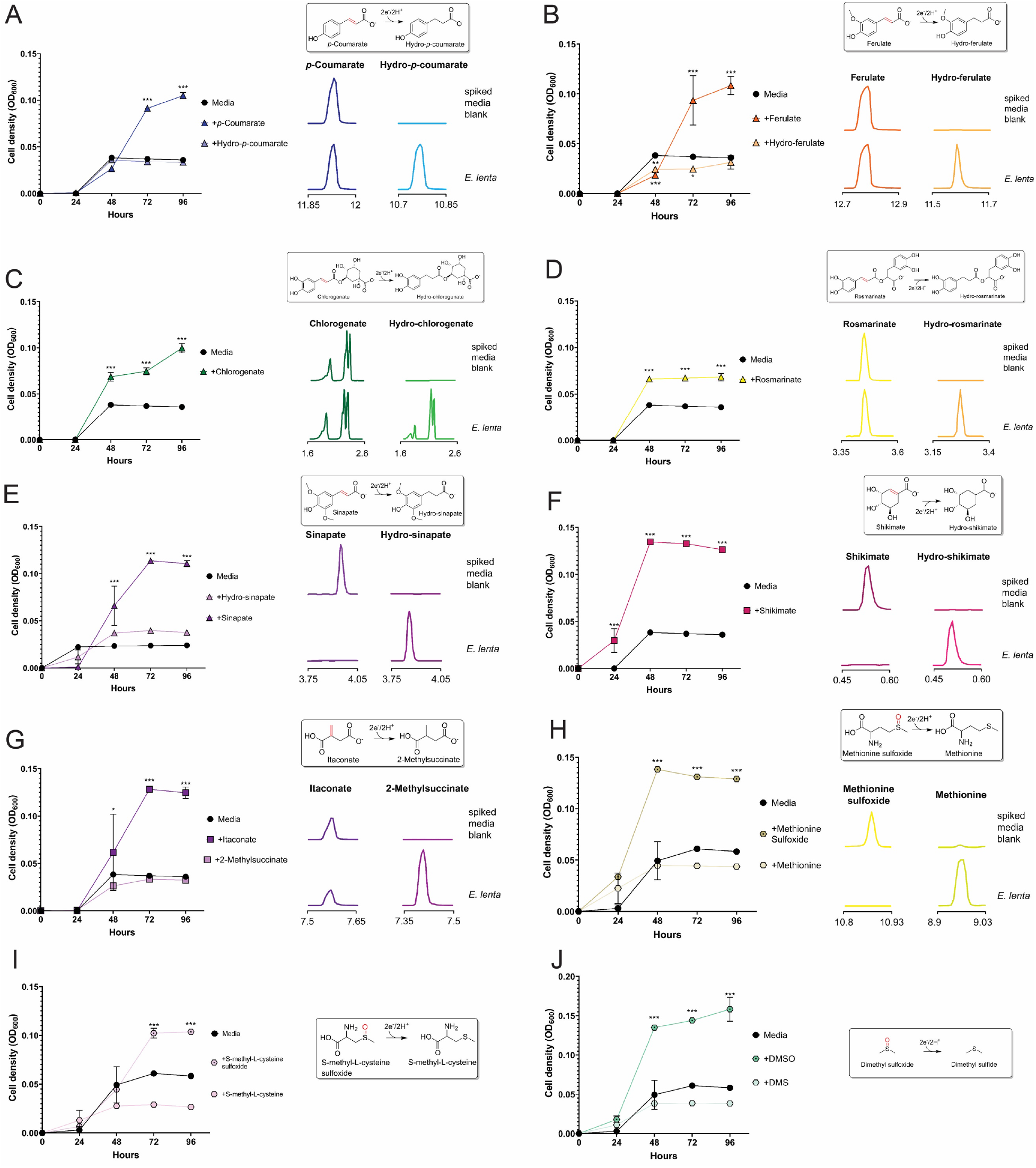
*E. lenta* uses multiple respiratory electron acceptors. *E. lenta* DSM2243 growth in formate-supplemented media provisioned with electron acceptors: (A) *p*-coumarate, (B) ferulate, (C) chlorogenate, (D) rosmarinate, (E) sinapate, (F) shikimate, (G) itaconate, (H) methionine sulfoxide, (I) S-methyl-L-cysteine sulfoxide, and (J) dimethyl sulfoxide. A ‘no electron acceptor’ condition and conditions with predicted reduction products are included as controls. Extracted ion chromatograms of peaks (matched to available authentic standards) in uninoculated and inoculated growth media. (A)-(D),(G)-(J) were measured by GC-MS; (E), (F) were measured by LC-MS; support for identification of hydro-shikimate in (F) is provided in **Supplementary Fig. 5**. Data are mean ±SD (n = 3 independent biological replicates). *p < 0.05, **p < 0.01, *** p < 0.001. Two-way ANOVA, multiple test vs media alone.

**Extended Data Figure 3.**
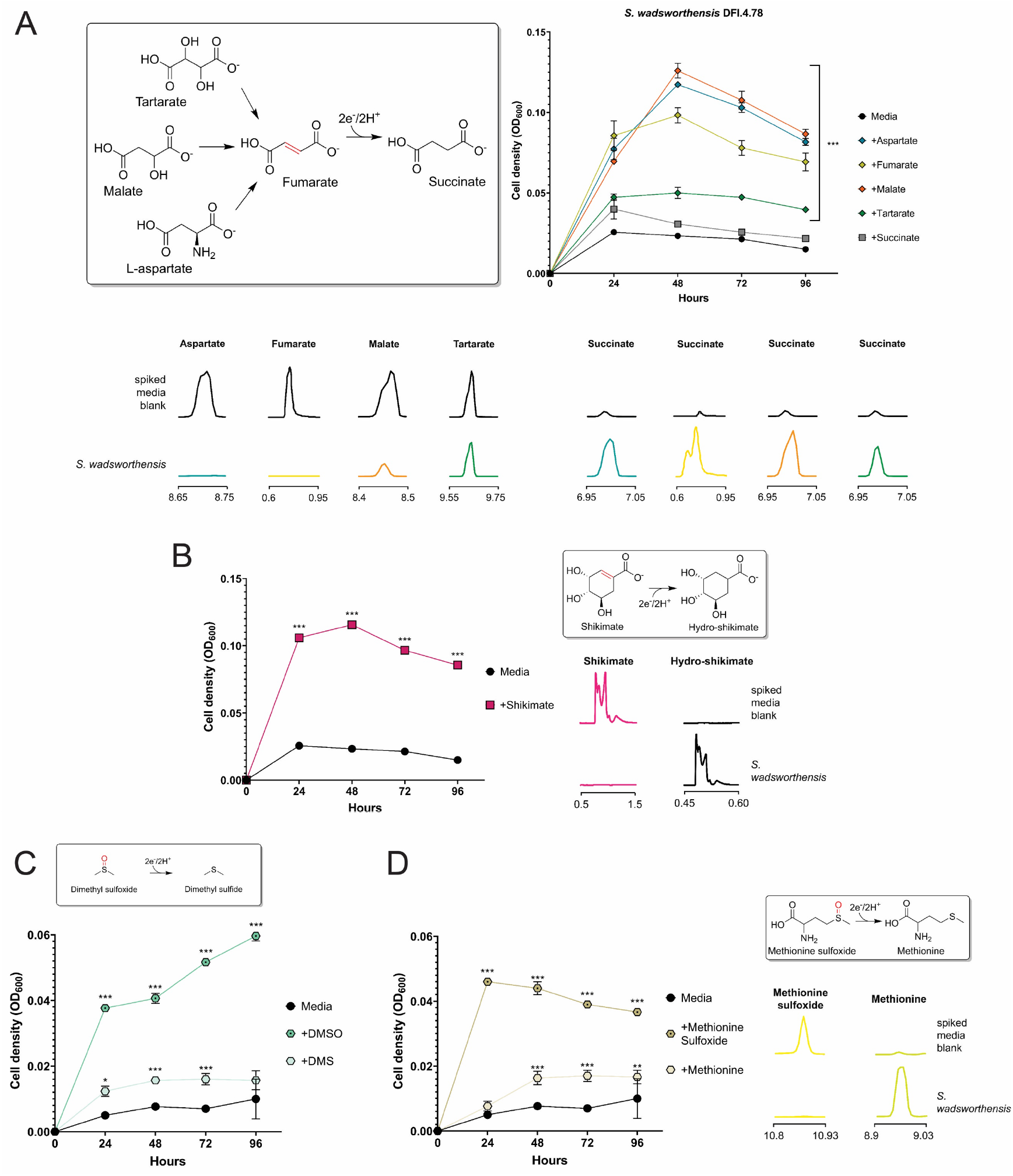
*S. wadsworthensis* growth-stimulating electron acceptors. *S. wadsworthensis* growth in formate-supplemented media provisioned with electron acceptors: (A) labeled C4-dicarboxylates, (B) shikimate, (C) dimethyl sulfoxide, and (D) methionine sulfoxide. A ‘no electron acceptor’ condition and conditions with predicted products are included as controls. Extracted ion chromatograms of peaks (matched to available authentic standards) in uninoculated and inoculated growth media. (A) Fumarate and its succinate reduction were measured by LC-MS, the rest of (A) and (D) were measured by GC-MS; (B) was measured by LC-MS; support for identification of hydro-shikimate in (B) is provided in **Supplementary Fig. 5**. Data are mean ±SD (n = 3 independent biological replicates). *p < 0.05, **p < 0.01, *** p < 0.001. Two-way ANOVA, multiple test vs media alone.

**Extended Data Figure 4.**
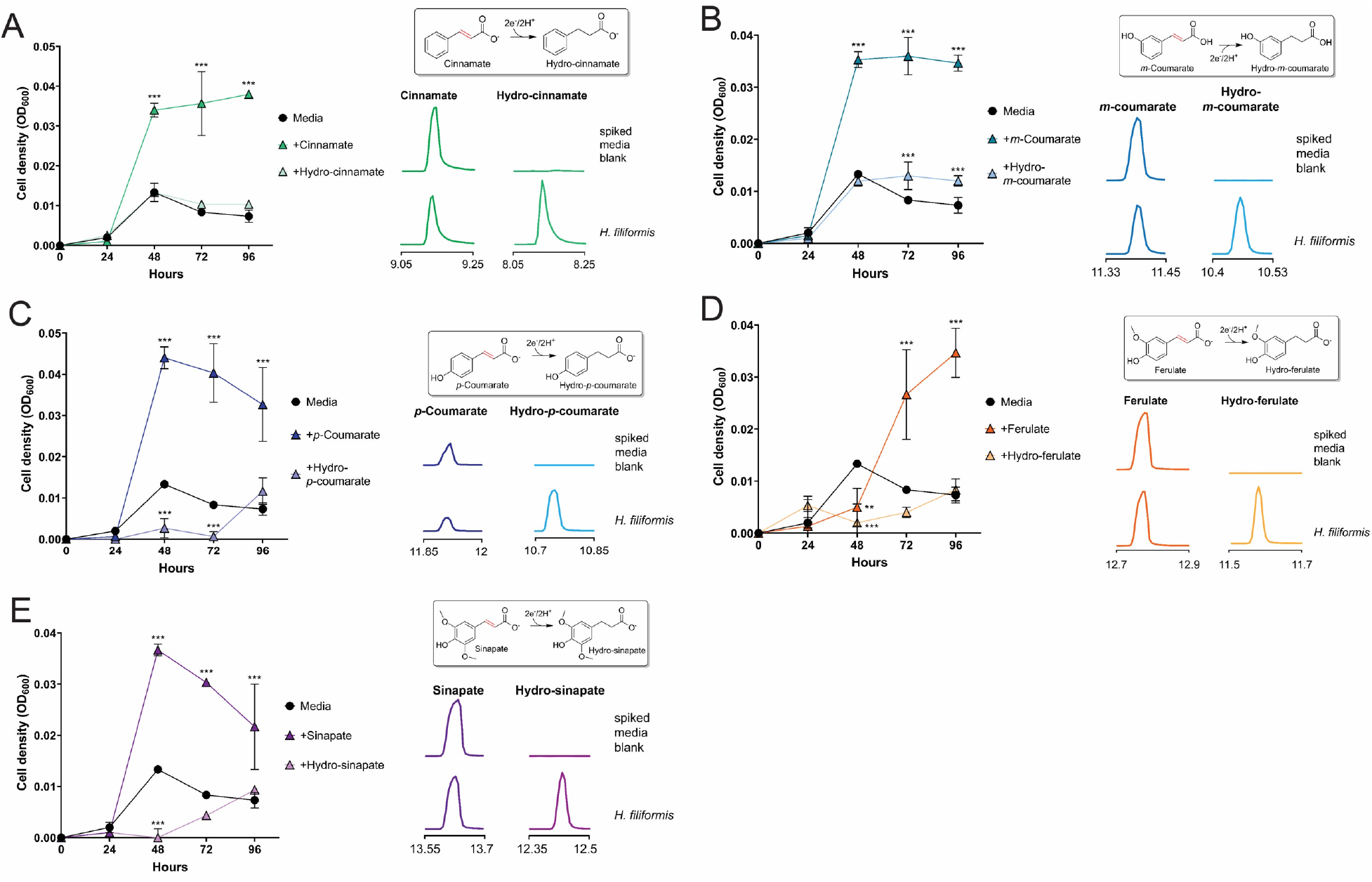
*H. filiformis* growth-stimulating electron acceptors. *H. filiformis* growth in media provisioned with electron acceptors: (A) cinnamate (B) *m*-coumarate, (C) *p*-coumarate (D) ferulate, and (E) sinapate. A ‘no electron acceptor’ condition and conditions with predicted products are included as controls. Extracted ion chromatograms of peaks (matched to available authentic standards) in uninoculated and inoculated growth media. Data are mean ±SD (n = 3 independent biological replicates). *p < 0.05, **p < 0.01, *** p < 0.001. Two-way ANOVA, multiple test vs media alone.

**Extended Data Figure 5.**
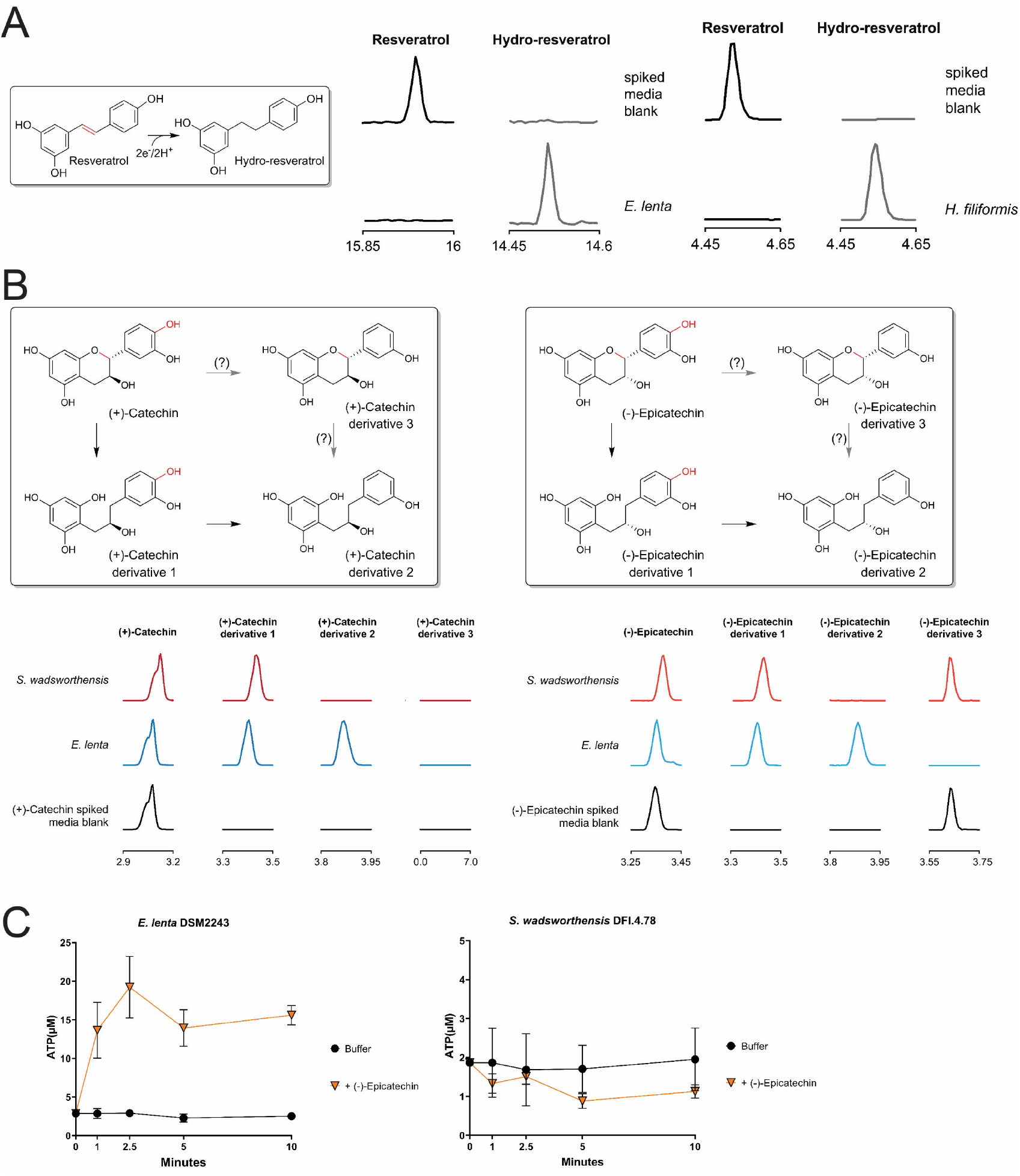
Electron acceptors not observed to stimulate growth. (A) GC-MS analysis of resveratrol-spiked media before and after *E. lenta* DSM2243 growth and LC-MS analysis of resveratrol-spiked media before and after *H. filiformis* DFI.9.20 growth. The low solubility of resveratrol hindered experiments to assess whether this electron acceptor supported respiratory growth. (B) Potential modification flow for (+)-catechin and (-)-epicatechin, and GC-MS of media spiked with either (+)-catechin or (-)-epicatechin before and after *E. lenta* DSM2243 or *S. wadsworthensis* DFI.4.78 growth. (C) Resting cell suspensions of *E. lenta* and *S. wadsworthensis* in buffer supplemented with 1mM formate assayed cellular ATP concentrations after incubation with buffer alone or (-)-epicatechin, data are mean ±SD (n = 3 technical replicates). Support for identification of catechin and epicatechin derivatives in (B) is provided in **Supplementary Fig. 2 & 3**.

**Extended Data Figure 6.**
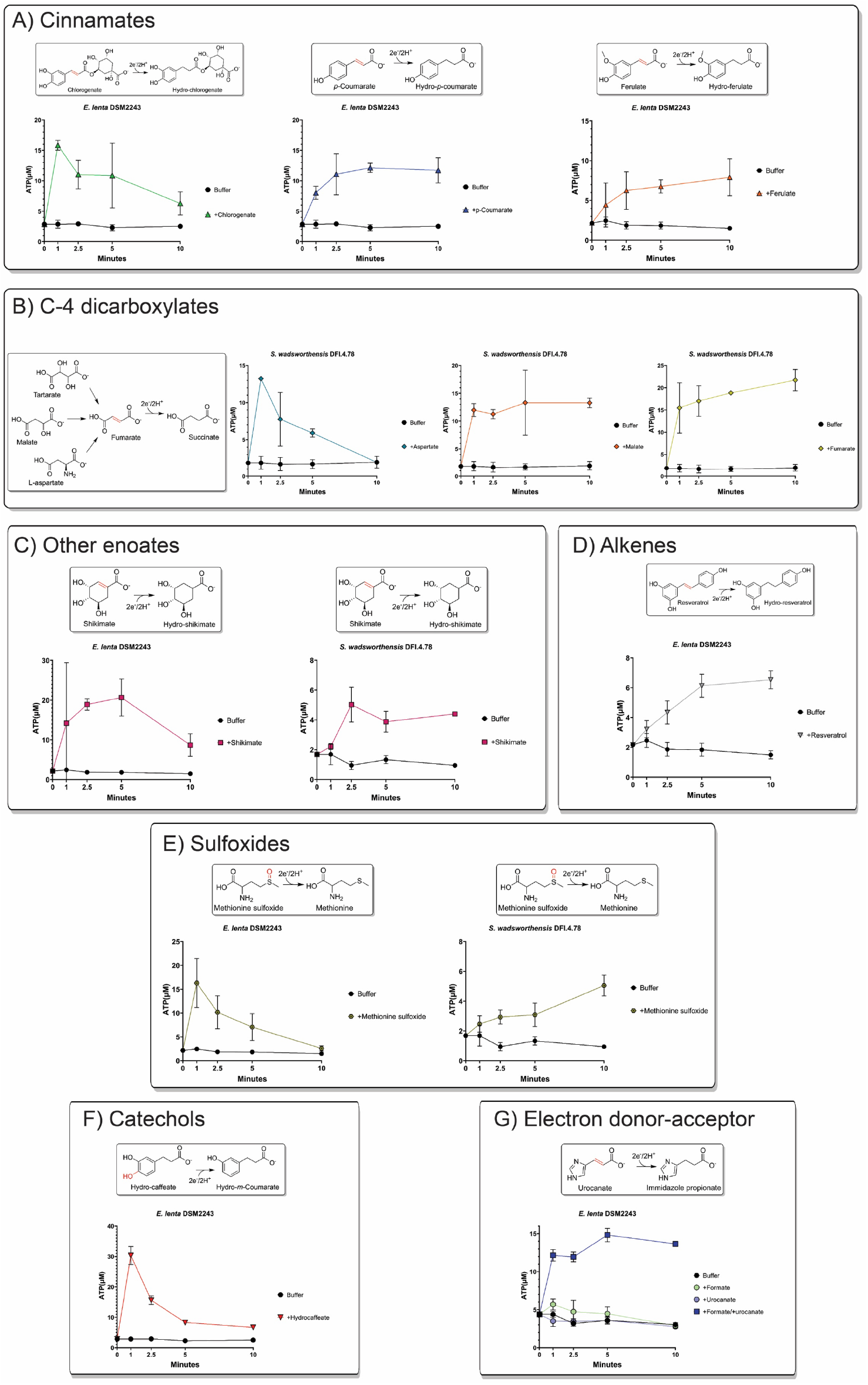
ATP generation facilitated by electron acceptors. Resting cell suspensions of *E. lenta* and *S. wadsworthensis* in buffer supplemented with 1mM formate assayed cellular ATP concentrations after incubation with different classes of electron acceptors, (A) cinnamates, (B) C4-dicarboxylates, (C) other enolates, (D) alkenes, (E) sulfoxides, (F) catechols, and (G) electron donor-acceptor combination dependence. Data are mean ±SD (n = 3 technical replicates).

**Extended Data Figure 7.**
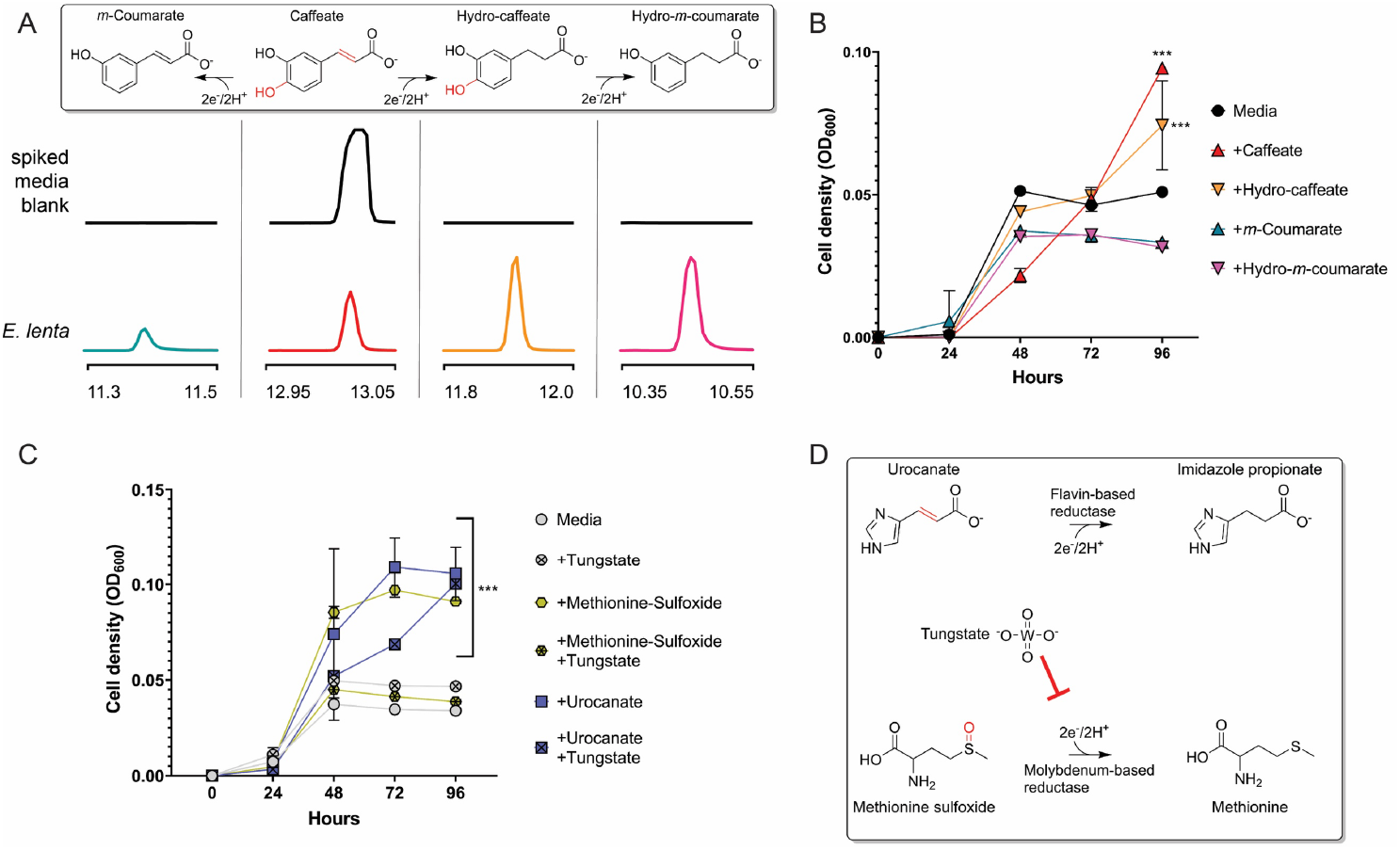
Caffeate utilization by *E. lenta* and tungstate inhibition of sulfoxide growth enhancement. (A) GC-MS analysis of supernatant collected from *E. lenta* DSM2243 grown in caffeate-spiked media. Extracted ion chromatograms of peaks (matched to authentic standards) in uninoculated and inoculated growth media. Proposed reaction pathways are shown, with peaks for each compound provided beneath its chemical structure. The previously characterized hydrocaffeate dehydroxylase may catalyze the observed dehydroxylation reactions. The observed caffeate reduction to hydrocaffeate provides evidence of a caffeate reductase, while the accumulation of *m*-coumarate suggests that this enzyme may specifically use caffeate. (B) *E. lenta* DSM2243 growth in media supplemented with formate and different cinnamates. The pattern of cinnamate-dependent growth enhancement supports the conclusions that: (1) dehydroxylation can support respiratory growth and (2) *m*-coumarate is a poor electron acceptor for *E. lenta*. (C) The effect of the molybdopterin reductase inhibitor, tungstate, on *E. lenta* DSM2243 growth. Media was supplemented with formate and the noted electron acceptor, with or without the addition of tungstate. (D) Reactions catalyzed by urocanate and sulfoxide reductases. Tungstate’s selective growth inhibition is consistent with sulfoxide, but not urocanate, reduction being catalyzed by a molybdopterin reductase. Data are mean ±SD (n = 3 independent biological replicates). *p < 0.05, **p < 0.01, *** p < 0.001. Two-way ANOVA, multiple test vs media alone.

**Extended Data Figure 8.**
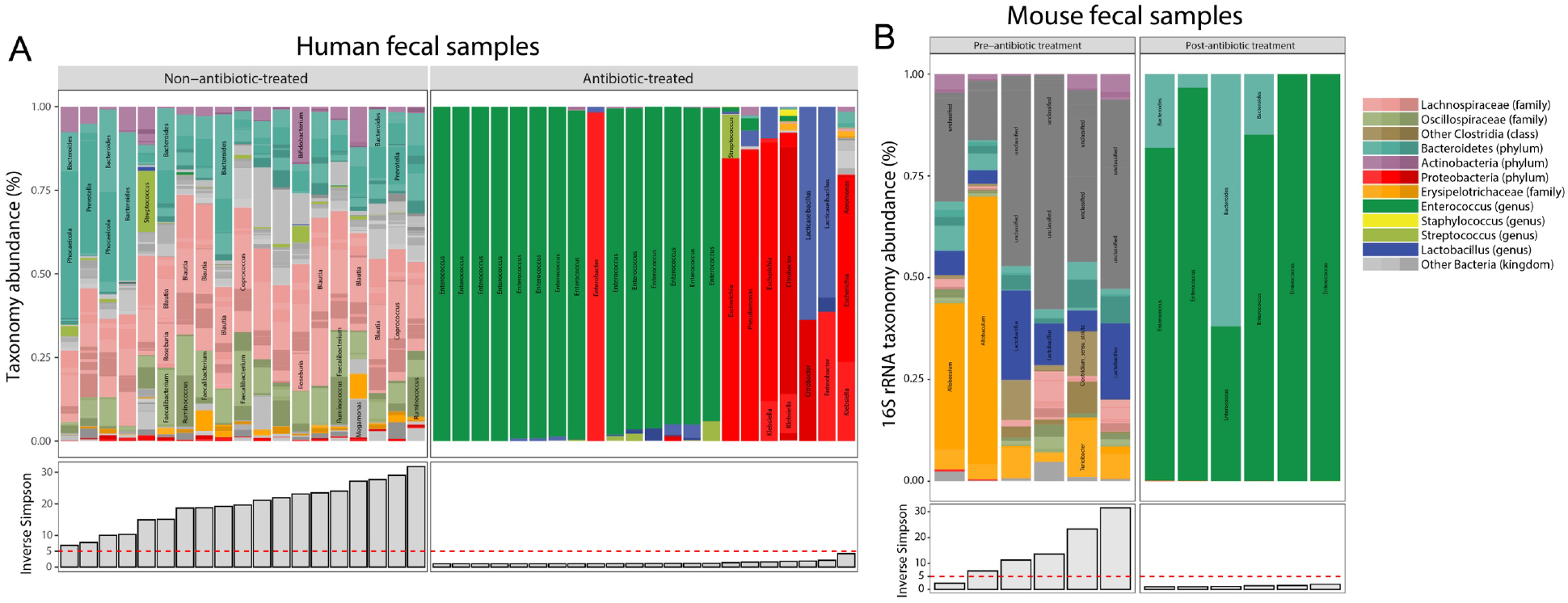
Microbiome composition of fecal samples used for metabolite measurements. (A) Taxonomy abundance in human fecal samples used for metabolomics analyses assessed by shotgun metagenomics. (B) Taxonomy abundance in mouse fecal samples used for metabolomics analyses assessed by 16S rRNA amplicon sequencing. The Inverse Simpson measure of microbiome diversity is also presented.

**Extended Data Figure 9.**
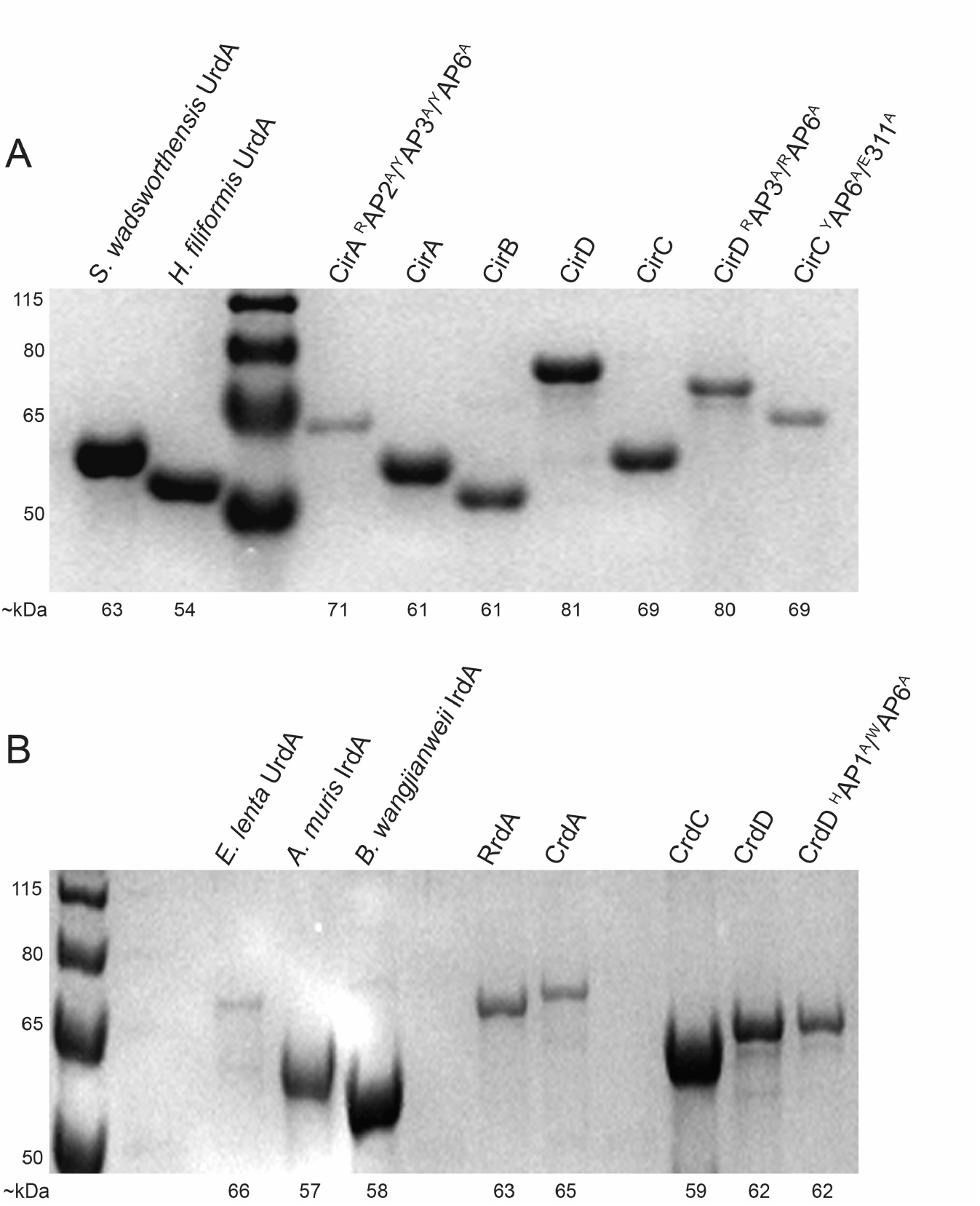
Expression of recombinant reductase enzymes. SDS-PAGE gels of reductase enzymes expressed and purified from *E. coli*. (A) *H. filiformis* enzymes, except where indicated. (B) *E. lenta* enzymes, except where indicated. Proteins run on a 12% acrylamide Bis-Tris gel in a MOPS running buffer, compared to PageRuler Plus prestained protein ladder (Thermo. #26619). Expected kDa for both ladder and proteins are labeled. Experiments (A, B) were repeated twice, with similar results.

**Extended Data Figure 10.**
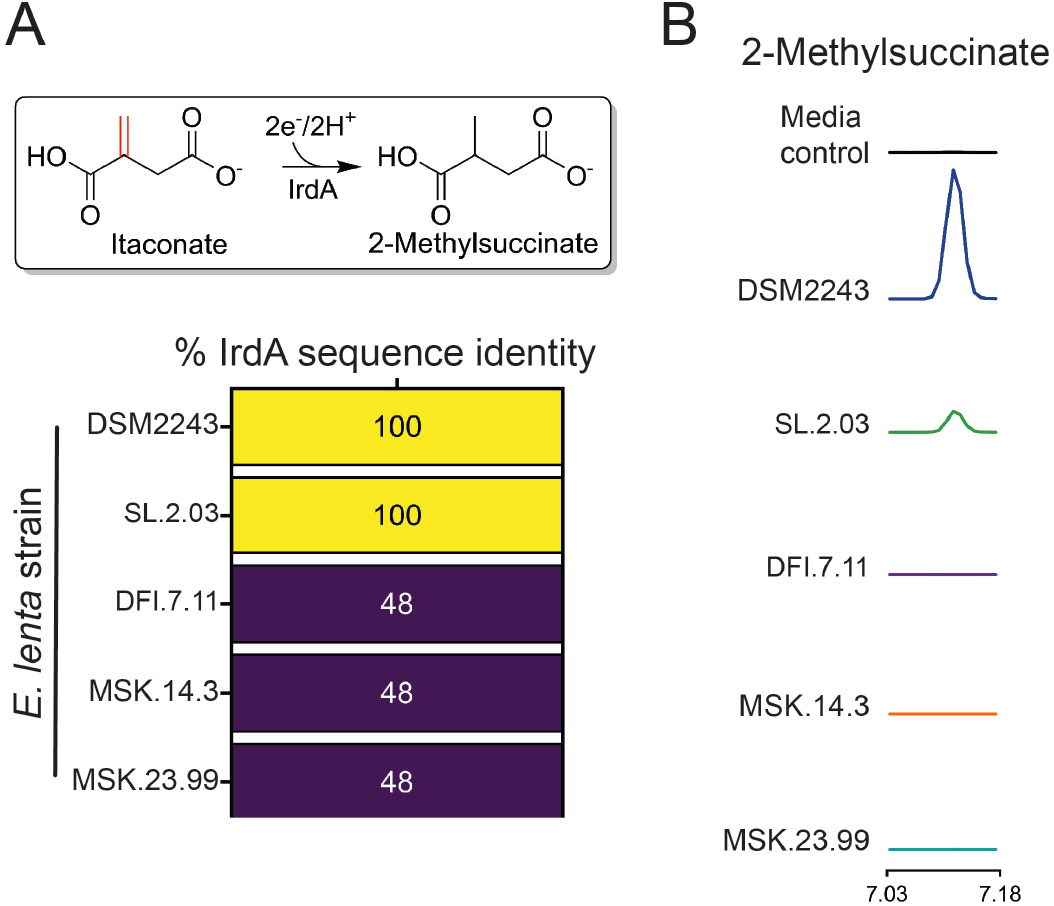
Presence of *irdA* predicts itaconate reductase activity of *E. lenta* strains. (A) Schematic diagram of itaconate reduction by IrdA, and sequence identity of the reductase with greatest similarity to IrdA is shown for indicated *E. lenta* strains strain. (B) Extracted ion chromatograms of authentic standard-matched methylsuccinate peaks from media collected after cultivation with the indicated strain.

## References

1. Moodie, A. D. & Ingledew, W. J. Microbial anaerobic respiration. Adv Microb Physiol 31, 225– 69 (1990).

2. Gibson, G. R., Macfarlane, G. T. & Cummings, J. H. Occurrence of sulphate-reducing bacteria in human faeces and the relationship of dissimilatory sulphate reduction to methanogenesis in the large gut. Journal of Applied Bacteriology 65, 103–111 (1988).

3. Smith, N. W., Shorten, P. R., Altermann, E. H., Roy, N. C. & McNabb, W. C. Hydrogen cross-feeders of the human gastrointestinal tract. Gut Microbes 10, 270–288 (2019).

4. Butler, N. L. et al. Bacteroides fragilis Maintains Concurrent Capability for Anaerobic and Nanaerobic Respiration. J Bacteriol 205, (2023).

5. Schubert, C. & Unden, G. C4-Dicarboxylates as Growth Substrates and Signaling Molecules for Commensal and Pathogenic Enteric Bacteria in Mammalian Intestine. J Bacteriol 204, e0054521 (2022).

6. Winter, S. E. et al. Gut inflammation provides a respiratory electron acceptor for Salmonella. Nature 467, 426–429 (2010).

7. Winter, S. E. et al. Host-derived nitrate boosts growth of E. coli in the inflamed gut. Science (1979) 339, 708–711 (2013).

8. Miller, B. M. et al. Anaerobic Respiration of NOX1-Derived Hydrogen Peroxide Licenses Bacterial Growth at the Colonic Surface. Cell Host Microbe 28, 789–797.e5 (2020).

9. Rekdal, V. M. et al. A widely distributed metalloenzyme class enables gut microbial metabolism of host-and diet-derived catechols. Elife 9, (2020).

10. Ravcheev, D. A. & Thiele, I. Systematic genomic analysis reveals the complementary aerobic and anaerobic respiration capacities of the human gut microbiota. Front Microbiol 5, 1–14 (2014).

11. Bilous, P. T., Cole, S. T., Anderson, W. F. & Weiner, J. H. Nucleotide sequence of the dmsABC operon encoding the anaerobic dimethylsulphoxide reductase of Escherichia coli. Mol Microbiol 2, 785–95 (1988).

12. Silvestro, A., Pommier, J., Pascal, M. C. & Giordano, G. The inducible trimethylamine N-oxide reductase of Escherichia coli K12: its localization and inducers. Biochim Biophys Acta 999, 208–16 (1989).

13. Heinzinger, N. K., Fujimoto, S. Y., Clark, M. A., Moreno, M. S. & Barrett, E. L. Sequence analysis of the phs operon in Salmonella typhimurium and the contribution of thiosulfate reduction to anaerobic energy metabolism. J Bacteriol 177, 2813–20 (1995).

14. Hensel, M., Hinsley, A. P., Nikolaus, T., Sawers, G. & Berks, B. C. The genetic basis of tetrathionate respiration in Salmonella typhimurium. Mol Microbiol 32, 275–87 (1999).

15. Cruz-García, C., Murray, A. E., Klappenbach, J. A., Stewart, V. & Tiedje, J. M. Respiratory nitrate ammonification by Shewanella oneidensis MR-1. J Bacteriol 189, 656–62 (2007).

16. Krafft, T. et al. Cloning and nucleotide sequence of the psrA gene of Wolinella succinogenes polysulphide reductase. Eur J Biochem 206, 503–10 (1992).

17. Saltikov, C. W. & Newman, D. K. Genetic identification of a respiratory arsenate reductase. Proc Natl Acad Sci U S A 100, 10983–8 (2003).

18. Krafft, T., Bowen, A., Theis, F. & Macy, J. M. Cloning and sequencing of the genes encoding the periplasmic-cytochrome B-containing selenate reductase of Thauera selenatis. DNA Seq 10, 365–77 (2000).

19. Bender, K. S. et al. Identification, characterization, and classification of genes encoding perchlorate reductase. J Bacteriol 187, 5090–6 (2005).

20. McPherson, M. J., Baron, A. J., Pappin, D. J. & Wootton, J. C. Respiratory nitrate reductase of Escherichia coli. Sequence identification of the large subunit gene. FEBS Lett 177, 260–4 (1984).

21. Lledó, B., Martínez-Espinosa, R. M., Marhuenda-Egea, F. C. & Bonete, M. J. Respiratory nitrate reductase from haloarchaeon Haloferax mediterranei: biochemical and genetic analysis. Biochim Biophys Acta 1674, 50–9 (2004).

22. Yamazaki, C. et al. A novel dimethylsulfoxide reductase family of molybdenum enzyme, Idr, is involved in iodate respiration by Pseudomonas sp. SCT. Environ Microbiol 22, 2196–2212 (2020).

23. Cole, S. T. Nucleotide sequence coding for the flavoprotein subunit of the fumarate reductase of Escherichia coli. Eur J Biochem 122, 479–84 (1982).

24. Light, S. H. et al. Extracellular electron transfer powers flavinylated extracellular reductases in Gram-positive bacteria. Proc Natl Acad Sci U S A 116, 26892–26899 (2019).

25. Bogachev, A. V., Bertsova, Y. V., Bloch, D. A. & Verkhovsky, M. I. Urocanate reductase: Identification of a novel anaerobic respiratory pathway in Shewanella oneidensis MR-1. Mol Microbiol 86, 1452–1463 (2012).

26. Speich, N. et al. Adenylylsulphate reductase from the sulphate-reducing archaeon Archaeoglobus fulgidus: cloning and characterization of the genes and comparison of the enzyme with other iron-sulphur flavoproteins. Microbiology (Reading) 140 **( Pt 6**, 1273–84 (1994).

27. Méheust, R., Huang, S., Rivera-Lugo, R., Banfield, J. F. & Light, S. H. Post-translational flavinylation is associated with diverse extracytosolic redox functionalities throughout bacterial life. Elife 10, 1–22 (2021).

28. Mikoulinskaia, O., Akimenko, V., Galouchko, A., Thauer, R. K. & Hedderich, R. Cytochrome c-dependent methacrylate reductase from Geobacter sulfurreducens AM-1. Eur J Biochem 263, 346–352 (1999).

29. Jardim-Messeder, D. et al. Fumarate reductase superfamily: A diverse group of enzymes whose evolution is correlated to the establishment of different metabolic pathways. Mitochondrion 34, 56–66 (2017).

30. Le, C. (Chip), Bae, M., Kiamehr, S. & Balskus, E. P. Emerging Chemical Diversity and Potential Applications of Enzymes in the DMSO Reductase Superfamily. Annu Rev Biochem 91, 475–504 (2022).

31. Almeida, A. et al. A unified catalog of 204,938 reference genomes from the human gut microbiome. Nat Biotechnol 39, 105–114 (2021).

32. Wu, S. et al. GMrepo: a database of curated and consistently annotated human gut metagenomes. Nucleic Acids Res 48, D545–D553 (2020).

33. Diaz-Gerevini, G. T. et al. Beneficial action of resveratrol: How and why? Nutrition 32, 174– 178 (2016).

34. Bentley, R. & Haslam, E. The Shikimate Pathway — A Metabolic Tree with Many Branche. Crit Rev Biochem Mol Biol 25, 307–384 (1990).

35. Michelucci, A. et al. Immune-responsive gene 1 protein links metabolism to immunity by catalyzing itaconic acid production. Proc Natl Acad Sci U S A 110, 7820–7825 (2013).

36. Dong, X. et al. Genetic manipulation of the human gut bacterium Eggerthella lenta reveals a widespread family of transcriptional regulators. Nat Commun 13, 7624 (2022).

37. Leys, D. et al. Structure and mechanism of the flavocytochrome c fumarate reductase of Shewanella putrefaciens MR-1. Nat Struct Biol 6, 1113–1117 (1999).

38. Pankhurst, K. L. et al. A proton delivery pathway in the soluble fumarate reductase from Shewanella frigidimarina. J Biol Chem 281, 20589–97 (2006).

39. Venskutonytė, R. et al. Structural characterization of the microbial enzyme urocanate reductase mediating imidazole propionate production. Nat Commun 12, 1347 (2021).

40. Heidelberg, J. F. et al. Genome sequence of the dissimilatory metal ion–reducing bacterium Shewanella oneidensis. Nat Biotechnol 20, 1118–1123 (2002).

41. Hau, H. H. & Gralnick, J. A. Ecology and Biotechnology of the Genus *Shewanella*. Annu Rev Microbiol 61, 237–258 (2007).

42. Ikeda, S. et al. Shewanella oneidensis MR-1 as a bacterial platform for electro-biotechnology. Essays Biochem 65, 355–364 (2021).

43. Koh, A. et al. Microbially Produced Imidazole Propionate Impairs Insulin Signaling through mTORC1. Cell 175, 947–961.e17 (2018).

44. Dodd, D. et al. A gut bacterial pathway metabolizes aromatic amino acids into nine circulating metabolites. Nature 551, 648–652 (2017).

45. Steed, A. L. et al. The microbial metabolite desaminotyrosine protects from influenza through type I interferon. Science (1979) 357, 498–502 (2017).

46. Haiser, H. J. et al. Predicting and Manipulating Cardiac Drug Inactivation by the Human Gut Bacterium Eggerthella lenta. Science (1979) 341, 295–298 (2013).

47. Koppel, N., Bisanz, J. E., Pandelia, M.-E., Turnbaugh, P. J. & Balskus, E. P. Discovery and characterization of a prevalent human gut bacterial enzyme sufficient for the inactivation of a family of plant toxins. Elife 7, 1–32 (2018).

48. Alexander, M. et al. Human gut bacterial metabolism drives Th17 activation and colitis. Cell Host Microbe 30, 17–30.e9 (2022).

49. Bess, E. N. et al. Genetic basis for the cooperative bioactivation of plant lignans by Eggerthella lenta and other human gut bacteria. Nat Microbiol 5, 56–66 (2020).

50. Maini Rekdal, V., Bess, E. N., Bisanz, J. E., Turnbaugh, P. J. & Balskus, E.P. Discovery and inhibition of an interspecies gut bacterial pathway for Levodopa metabolism. Science 364, (2019).

51. Sasikaran, J., Ziemski, M., Zadora, P. K., Fleig, A. & Berg, I. A. Bacterial itaconate degradation promotes pathogenicity. Nat Chem Biol 10, 371–7 (2014).

52. Wang, H. et al. An essential bifunctional enzyme in Mycobacterium tuberculosis for itaconate dissimilation and leucine catabolism. Proc Natl Acad Sci U S A 116, 15907–15913 (2019).

53. Zhang, T., Hasegawa, Y. & Waldor, M. K. A bile metabolite atlas reveals infection-triggered interorgan mediators of intestinal homeostasis and defense. bioRxiv 2023.03.04.531105 (2023) doi:10.1101/2023.03.04.531105.

54. Almeida, A. et al. A unified catalog of 204,938 reference genomes from the human gut microbiome. Nat Biotechnol 39, 105–114 (2021).

55. Hyatt, D. et al. Prodigal: Prokaryotic gene recognition and translation initiation site identification. BMC Bioinformatics (2010) doi:10.1186/1471-2105-11-119.

56. Mistry, J. et al. Pfam: The protein families database in 2021. Nucleic Acids Res 49, D412– D419 (2021).

57. Eddy, S. R. Profile hidden Markov models. Bioinformatics Preprint at 10.1093/bioinformatics/14.9.755 (1998).

58. Almagro Armenteros, J. J., et al. SignalP 5.0 improves signal peptide predictions using deep neural networks. Nat Biotechnol 37, 420–423 (2019).

59. Katoh, K. & Standley, D. M. A simple method to control over-alignment in the MAFFT multiple sequence alignment program. Bioinformatics (2016) doi:10.1093/bioinformatics/btw108.

60. Capella-Gutiérrez, S., Silla-Martínez, J. M. & Gabaldón, T. trimAl: A tool for automated alignment trimming in large-scale phylogenetic analyses. Bioinformatics (2009) doi:10.1093/bioinformatics/btp348.

61. Nguyen, L. T., Schmidt, H. A., Von Haeseler, A. & Minh, B. Q. IQ-TREE: A fast and effective stochastic algorithm for estimating maximum-likelihood phylogenies. Mol Biol Evol (2015) doi:10.1093/molbev/msu300.

62. Kalyaanamoorthy, S., Minh, B. Q., Wong, T. K. F., Von Haeseler, A. & Jermiin, L. S. ModelFinder: Fast model selection for accurate phylogenetic estimates. Nat Methods (2017) doi:10.1038/nmeth.4285.

63. Hoang, D. T., Chernomor, O., Von Haeseler, A., Minh, B. Q. & Vinh, L. S. UFBoot2: Improving the ultrafast bootstrap approximation. Mol Biol Evol (2018) doi:10.1093/molbev/msx281.

64. Delmont, T. O. & Eren, A. M. Linking pangenomes and metagenomes: the Prochlorococcus metapangenome. PeerJ 6, e4320 (2018).

65. Altschul, S. F., Gish, W., Miller, W., Myers, E. W. & Lipman, D. J. Basic local alignment search tool. J Mol Biol 215, 403–410 (1990).

66. García-Villalba, R. et al. Metabolism of different dietary phenolic compounds by the urolithin-producing human-gut bacteria Gordonibacter urolithinfaciens and Ellagibacter isourolithinifaciens. Food Funct 11, 7012–7022 (2020).

67. Sumner, L. W. et al. Proposed minimum reporting standards for chemical analysis. Metabolomics 3, 211–221 (2007).

68. Odenwald, M. A. et al. Bifidobacteria metabolize lactulose to optimize gut metabolites and prevent systemic infection in patients with liver disease. Nat Microbiol 2058-5276 (2023) doi.10.1038/s41564-023-01493-w.

69. Bolger, A. M., Lohse, M. & Usadel, B. Trimmomatic: a flexible trimmer for Illumina sequence data. Bioinformatics 30, 2114–2120 (2014).

70. Blanco-Miguez, A. et al. Extending and improving metagenomic taxonomic profiling with uncharacterized species with MetaPhlAn 4. bioRxiv 2022.08.22.504593 (2022) doi:10.1101/2022.08.22.504593.

71. Langmead, B. & Salzberg, S. L. Fast gapped-read alignment with Bowtie 2. Nat Methods 9, 357–359 (2012).

72. Danecek, P. et al. Twelve years of SAMtools and BCFtools. Gigascience 10, (2021).

73. Liao, Y., Smyth, G. K. & Shi, W. featureCounts: an efficient general purpose program for assigning sequence reads to genomic features. Bioinformatics 30, 923–930 (2014).

74. Love, M. I., Huber, W. & Anders, S. Moderated estimation of fold change and dispersion for RNA-seq data with DESeq2. Genome Biol 15, 550 (2014).

75. Team, R. C. R: A language and environment for statistical computing. R Foundation for Statistical Computing, Vienna, Austria. http://www.R-project.org/ (2016).

76. Zhang, Y., Parmigiani, G. & Johnson, W. E. ComBat-seq: batch effect adjustment for RNA-seq count data. NAR Genom Bioinform 2, (2020).

77. Zhu, A., Ibrahim, J. G. & Love, M. I. Heavy-tailed prior distributions for sequence count data: removing the noise and preserving large differences. Bioinformatics 35, 2084–2092 (2019).

78. Plumb, R. S. et al. UPLC/MS(E); a new approach for generating molecular fragment information for biomarker structure elucidation. Rapid Commun Mass Spectrom 20, 1989–94 (2006).

79. Shliaha, P. V, Bond, N. J., Gatto, L. & Lilley, K. S. Effects of traveling wave ion mobility separation on data independent acquisition in proteomics studies. J Proteome Res 12, 2323– 39 (2013).

80. Helm, D. et al. Ion mobility tandem mass spectrometry enhances performance of bottom-up proteomics. Mol Cell Proteomics 13, 3709–15 (2014).

81. Distler, U. et al. Drift time-specific collision energies enable deep-coverage data-independent acquisition proteomics. Nat Methods 11, 167–70 (2014).

82. Distler, U., Kuharev, J., Navarro, P. & Tenzer, S. Label-free quantification in ion mobility-enhanced data-independent acquisition proteomics. Nat Protoc 11, 795–812 (2016).

83. Sayers, E. W. et al. Database resources of the national center for biotechnology information. Nucleic Acids Res 50, D20–D26 (2022).

84. Rivera-Lugo, R. et al. Distinct Energy-Coupling Factor Transporter Subunits Enable Flavin Acquisition and Extracytosolic Trafficking for Extracellular Electron Transfer in Listeria monocytogenes. mBio 14, e0308522 (2023).

85. Eren, A. M. et al. Community-led, integrated, reproducible multi-omics with anvi’o. Nat Microbiol 6, 3–6 (2020).

86. Eddy, S. R. Accelerated Profile HMM Searches. PLoS Comput Biol 7, e1002195 (2011).

87. Steinegger, M. & Söding, J. MMseqs2 enables sensitive protein sequence searching for the analysis of massive data sets. Nature Biotechnology Preprint at 10.1038/nbt.3988 (2017).

88. Edgar, R. C. MUSCLE: multiple sequence alignment with high accuracy and high throughput. Nucleic Acids Res 32, 1792–1797 (2004).

89. Minh, B. Q. et al. IQ-TREE 2: New Models and Efficient Methods for Phylogenetic Inference in the Genomic Era. Mol Biol Evol 37, 1530–1534 (2020).

90. Jumper, J. et al. Highly accurate protein structure prediction with AlphaFold. Nature 596, 583– 589 (2021).

91. Waterhouse, A. M., Procter, J. B., Martin, D. M. A., Clamp, M. & Barton, G. J. Jalview Version 2--a multiple sequence alignment editor and analysis workbench. Bioinformatics 25, 1189–1191 (2009).

